# Gut Microbiome Signatures Linked to HIV-1 Reservoir Size and Viremia Control

**DOI:** 10.1101/2021.10.03.462590

**Authors:** Alessandra Borgognone, Marc Noguera-Julian, Bruna Oriol, Laura Noël-Romas, Marta Ruiz-Riol, Yolanda Guillén, Mariona Parera, Maria Casadellà, Clara Duran, Maria C. Puertas, Francesc Català-Moll, Marlon De Leon, Samantha Knodel, Kenzie Birse, Christian Manzardo, Jose M. Miró, Bonaventura Clotet, Javier Martinez-Picado, José Moltó, Beatriz Mothe, Adam Burgener, Christian Brander, Roger Paredes, the BCN02 Study Group

## Abstract

**Background:** The potential role of the gut microbiome as a predictor of immune-mediated HIV-1 control in the absence of antiretroviral therapy (ART) is still unknown. In the BCN02 clinical trial, which combined the MVA.HIVconsv immunogen with the latency-reversing agent romidepsin in early-ART treated HIV-1 infected individuals, 23% (3/13) of participants showed sustained low-levels of plasma viremia during 32 weeks of a monitored ART pause (MAP). Here, we present a multi-omics analysis to identify compositional and functional gut microbiome patterns associated with HIV-1 control in the BCN02 trial.

**Results:** Viremic controllers during the MAP (controllers) exhibited higher *Bacteroidales/Clostridiales* ratio and lower microbial gene richness before vaccination and throughout the study intervention when compared to non-controllers. Longitudinal assessment indicated that the gut microbiome of controllers was enriched in pro-inflammatory bacteria and depleted in butyrate-producing bacteria and methanogenic archaea. Functional profiling also showed that metabolic pathways, including methanogenesis and carbohydrate biosynthesis, were significantly decreased in controllers. Fecal metaproteome analyses confirmed that baseline functional differences were mainly driven by *Clostridial*es. Participants with high baseline *Bacteroidales/Clostridiales* ratio had increased pre-existing immune activation-related transcripts. The *Bacteroidales/Clostridiales* ratio as well as host immune-activation signatures inversely correlated with HIV-1 reservoir size.

**Conclusions:** This proof-of-concept study suggests the *Bacteroidales/Clostridiales* ratio as a novel gut microbiome signature associated with HIV-1 reservoir size and immune-mediated viral control after ART interruption.

## Background

A major obstacle to HIV-1 cure is the persistence of viral reservoirs. This mainly refers to latently-infected cells carrying transcriptionally-silent, replication-competent viruses which evade antiretroviral therapy (ART) as well as immune-mediated clearance[1–3]. The immune system is generally unable to contain HIV-1 replication in the absence of ART[4]. However, up to 10-20% of subjects that initiate ART within first weeks after HIV-1 acquisition may temporarily achieve HIV-1 viremia suppression after ART interruption (ATI)[5]. Understanding the mechanisms behind immune-mediated viremia control after ATI is key to progress towards a functional HIV cure. Broader and higher-magnitude CTL (cytotoxic T-lymphocyte) responses against less diverse HIV-1 epitopes[6,7] in the context of favorable HLA class I genotypes[8] and smaller HIV-1 reservoir size[9] have all been related to such post-treatment HIV-1 control.

There is indirect evidence that the gut microbiome might also contribute to immune-mediated control of HIV-1 replication[10,11]. Vaccine-induced gut microbiome alterations, consisting in lower bacterial diversity and negative correlation between richness and CD14^+^DR^-^ monocytes in colorectal intraepithelial lymphocytes, have been recently associated with HIV/SIV (SHIV) protection in a non-human primate challenge study after mucosal vaccination with HIV/SIV peptides, modified vaccinia Ankara–SIV and HIV-gp120–CD4 fusion protein plus adjuvants through the oral route[12]. In the HVTN 096 trial[13], where the impact of the gut microbiota on HIV-specific immune response to a DNA-prime, poxvirus-boost strategy in human adults was assessed, baseline and vaccine-induced gp41-reactive IgG titers were associated with different microbiota community structures, in terms of richness and composition[14]. In particular, co-occurring bacterial groups, such as *Ruminococcaceae, Peptoniphilaceae*, and *Bacteroidaceae*, were associated with vaccine-induced IgG response and inversely correlated with pre-existing gp41 binding IgG antibodies, suggesting that the microbiome may influence the immune response and vaccine immunogenicity[15]. Further evidence emerged from other studies in typhoid Ty21[16], rotavirus[17] and oral polio virus, tetanus-toxoid, bacillus Calmette-Guérin and hepatitis B immunization strategies[18], in which specific gut microbiome signatures (*Bifidobacterium, Streptococcus bovis* and *Clostridiales*, respectively) positively correlated with vaccine-induced immune response. In the absence of immune correlates of viral control, HIV cure trials usually incorporate an ART interruption phase to address the efficacy of a therapeutic intervention[19]. Data on the role of gut microbiome composition in the responsiveness to a curative strategy and the relationship with viral control after ART interruption are lacking. The BCN02 study[20] was a single-arm, proof of concept “kick & kill” clinical trial evaluating the safety and the *in vivo* effects of the histone deacetylase inhibitor romidepsin given as a latency reversing agent[21] in combination with a therapeutic HIV vaccine (MVA.HIVconsv) in a group of early-ART treated HIV-1-infected individuals[22,23]. During a monitored ART interruption (MAP), 23% of individuals showed sustained viremia control up to 32 weeks of follow-up.

Here, we aimed to identify salient compositional and functional gut microbiome patterns associated with control of HIV-1 viremia after ART interruption in the “kick & kill” strategy used in the BCN02 study.

## Materials and Methods

### Study design

This was a sub-study derived from the BCN02 clinical trial (NCT02616874). The BCN02 was a multicenter, open-label, single-arm, phase I, proof of concept clinical trial in which 15 HIV-1-infected individuals with sustained viral suppression who started ART within the first six months after HIV transmission were enrolled to evaluate the safety, tolerability, immunogenicity and effect on the viral reservoir of a kick&kill strategy consisting of the combination of HIVconsv vaccines with romidepsin[20] (Additional Figures: Figure S1a). Fifteen individuals enrolled in the BCN02 trial (procedures for recruitment and eligibility criteria are detailed elsewhere[20]) were immunized with a first dose of MVA.HIVconsv (MVA1, 2 × 10^8^ pfu intramuscularly), followed by three weekly-doses of romidepsin (RMD_1-2-3_, 5 mg/m^2^ BSA intravenously) and a second boost of MVA.HIVconsv (MVA2, 2 × 10^8^ pfu intramuscularly) four weeks after the last RMD_3_ infusion. To assess the ability for viral control after ART interruption, participants underwent a monitored antiviral pause (MAP), 8 weeks after the second vaccination (MVA2), for a maximum of 32 weeks or until any ART resumption criteria were met (plasma viral load > 2,000 copies/ml, CD4+ cell counts < 500 cells/mm3 and/or development of clinical symptoms related to an acute retroviral syndrome[20]). The study was conducted between February 2016 and October 2017 at two HIV-1 units from university hospitals in Barcelona (Hospital Germans Trias i Pujol and Hospital Clínic) and a community center (BCN-Checkpoint, Barcelona). The microbiome sub-study concept, design and patient information were reviewed and approved by the institutional ethical review board of the participating institutions (Reference Nr AC-15-108-R) and by the Spanish Regulatory Authorities (EudraCT 2015-002300-84). Written informed consent was provided by all study participants in accordance to the principles expressed in the Declaration of Helsinki and local personal data protection law (LOPD 15/1999).

### Sample disposition and data analysis

Fourteen participants from the BCN02 trial consented to participate in the BCN02-microbiome study, one was excluded due to a protocol violation during MAP and thirteen were included for multi-omics analyses. Twelve from the thirteen participants that finalized the “kick & kill” intervention completed the MAP phase (n=3 controllers and n=9 non-controllers) and one subject (B07) did not enter the MAP period due to immune futility pre-defined criteria and absence of protective HLA class I protective alleles associated with natural HIV-1 control (Additional Figures: Figure S1b). Based on the gut microbiome similarity with non-controllers at study entry and over the “kick & kill” intervention, the participant B07 was included in the non-controller arm to increase the statistical power in this microbiome sub-study.

Fecal specimens were longitudinally collected at BCN02 during the intervention period at study entry (pre-Vax), 1 week after 1^st^ vaccination (MVA1), 1 week after RMD_3_ (RMD) and 4 weeks after 2^nd^ vaccination (MVA2). Samples were also collected over the MAP period (from 4 to 34 weeks after ART interruption) and 24 weeks after ART resumption (Additional Figures: Figure S1a). All samples were processed for shotgun metagenomics analysis. Taxonomical classification, microbial gene content and functional profiling were inferred using Metaphlan2[24], IGC reference catalog[25] and HUMAnN2[26], respectively. Sequencing analysis and quality control of metagenomics data are provided in the Additional Results section (Additional Text). To facilitate the interpretation, longitudinal time points were schematically grouped into three phases (Additional Figures: Figure S2a). Fecal material, peripheral blood mononuclear cells (PBMC) and plasma samples were also sampled at baseline to assess fecal metaproteome, host transcriptome profiles and soluble inflammation biomarkers, respectively (Additional Figures: Figure S2b). Microbial proteins from fecal samples were measured by mass spectrometry and protein identification performed using Mascot search engine (v2.4, Matrix Science) and Scaffold Q+ software (v4.9.0, Proteome Software)[27]. PBMC transcriptomes were evaluated using RNA-sequencing and sequence reads aligned to the human reference genome by STAR v2.5.3a[28]. Read counts estimation was inferred using RSEM v1.3.0[29] and differential expression analysis performed by DESeq2[30]. Plasma proteins were estimated using the Proximity Extension Assay based on the Olink Inflammation Panel[31]. Correlations between ‘omic’ datasets were computed using Spearman’s correlation coefficients and integrative multi-omics analysis was assessed based on the mixOmics R package[32]. A detailed description of wet-lab procedures, bioinformatic methods and statistical analysis of metagenome, metaproteome, transcriptome, soluble plasma markers and multi-omics data is available in the Additional Methods section (Additional Text).

## Results

### Patient characteristics

In this microbiome sub-study, we evaluated 13 participants of the BCN02 study. Three had sustained plasma HIV-1 viremia (<2,000 copies/ml) during 32 weeks of MAP (viremic controllers), whereas 9 developed HIV-1 RNA rebound (>2,000 copies/ml) during MAP (non-controllers). One additional subject (B07) did not qualify for MAP due to pre-specified immune futility criteria and absence of protective HLA alleles, and therefore, was also considered a non-controller in this microbiome study. (Additional Figures: Figure S1b). Study participants were predominantly MSM (92%) of Caucasian ethnicity (92%), with median age of 42 years and median body mass index of 22.9 kg/m^2^ (Table 1). Median baseline CD4^+^ T-cell counts were 728 (416-1408) cells/ mm^3^ and median CD4/CD8 T-cell ratio was 1.4 (0.97-1.9). All subjects had been on integrase strand-transfer inhibitor-based triple ART for >3 years, begun during the first 3 months after HIV-1 infection. Median baseline HIV-1 proviral DNA was 140 copies/10^6^ CD4^+^ T-cells, being numerically lower in controllers than in non-controllers (65 *vs* 165 copies/10^6^ CD4^+^ T-cells, *p*=0.29).

**Table 1.** Study participant demographics and clinical characteristics.

| Variable | All participants<br>(n=13) | non-controllers<br>(n=10) | controllers<br>(n=3) |
| --- | --- | --- | --- |
| **Demographics** |  |  |  |
| Sex (M/F), *n* | 12/1 | 9/1 | 3/0 |
| Risk group (MSM/HTS), *n* | 12/1 | 9/1 | 3/0 |
| Ethnic group (Caucasian/Latin), *n* | 12/1 | 9/1 | 3/0 |
| Age (years) | 42 (39-47) | 43 (39-47) | 34 (33-38) |
| BMI (Kg/m<sup>2</sup>) | 22.9 (20.9-24) | 22.3 (21.1-23.4) | 24.3 (22.2-25) |
| **Treatment and clinical characteristics** |  |  |  |
| ART regimen, *n*<br>(TDF_FTC_RAL/ABC_3TC_RAL/ABC_3TC_DTG) | 2/9/2 | 2/6/2 | 0/3/0 |
| Viral reservoir (HIV-1 DNA cp/10<sup>6</sup> CD4<sup>+</sup> T-cells) | 140 (65-361) | 165 (76.2-415.7) | 65 (62.5-116.5) |
| CD4<sup>+</sup> T-cell (cells/mm<sup>3</sup>) | 728 (648-1182) | 839 (581.8-1293.8) | 657 (652.5-814) |
| CD4<sup>+</sup> T-cell (%) | 42.9 (42.2-49.3) | 43.4 (42.3-48.1) | 42.2 (38.4-48.1) |
| CD4/CD8 T-cell counts ratio | 1.4 (1.2-1.6) | 1.4 (1.2-1.5) | 1.3 (1.1-1.6) |

Continuous data are presented using median, 25% and 75% interquartile range, unless otherwise described.
M, male; F, female; MSM, men who have sex with men; HTS, heterosexual; BMI, body mass index; ART, antiretroviral therapy; cp, copies; TDF, Tenofovir Disoproxil Fumarate; FTC, Emtricitabine; RAL, Raltegravir; ABC, Abacavir; 3TC, Lamivudine; DTG, Dolutegravir. No statistically significant differences were observed ( $p \leq 0.05$ ; Wilcoxon rank-sum test

### Baseline gut-associated *Bacteroidales / Clostridiales* ratio discriminates between viremic controllers and non-controllers

Viremic controllers had significantly higher *Bacteroidales* levels than non-controllers at study entry (pre-Vax) (*p* = 0.007) and during all the intervention phase (after the first MVA dose: MVA1 *p* = 0.049, after the three romidepsin doses: RMD *p* = 0.049 and after the second MVA dose: MVA2, *p* = 0.014) (Fig. 1a, Additional Figures: Figures S3-S4) as well as lower *Clostridiales* (*p* = 0.014) levels before vaccination (Fig. 1b, Additional Figures: Figures S3-S4). The *Bacteroidales/Clostridiales* ratio remained significantly higher in controllers throughout the intervention (pre-Vax, *p* = 0.007 and MVA2, *p* = 0.028) (Fig. 1c). In addition, non-controllers were enriched in *Erysipelotrichales* and *Coriobacteriales* (Additional Figures: Figure S4a) and showed significantly higher *Methanobacteriales* levels (Additional Figures: Figure S4b). More detailed analyses at lower taxonomic levels using the LEfSe algorithm (Additional Figures: Figure S5) showed that controllers were mainly enriched in *Prevotella copri*, as well as in *Haemophilus pittmaniae* and *Streptococcus parasanguinis*. In comparison, non-controllers were enriched in the *Clostridiales* species *Eubacterium rectale and siraeum, Subdoligranulum spp. Coprococcus*, and *Dorea longicatena*, as well as in *Collinsella aerofaciens* and *Methanobrevibacter spp*. A longitudinal analysis using the ‘feature-volatility’ function from qiime2 (Additional Figures: Figure S6) showed that such differences were sustained over the whole intervention period. All 9 non-controllers analyzed here resumed ART by week 4 after the MAP initiation, whereas the 3 controllers remained off ART for at least 28 weeks and up to 32 weeks. During the MAP, *Bacteroidales* showed an initial increase up to week 4 followed by a reduction by weeks 8-12 in controllers (Fig. 1a). Inversely, *Clostridiales* levels increased by weeks 8-12 and remained stable thereafter (Fig. 1b) (no statistical support provided during MAP). No significant differences in bacterial composition were found following ART resumption; however, there was a limited sample availability at ART resumption phase (Figs. 1a-c).

**Fig 1.**
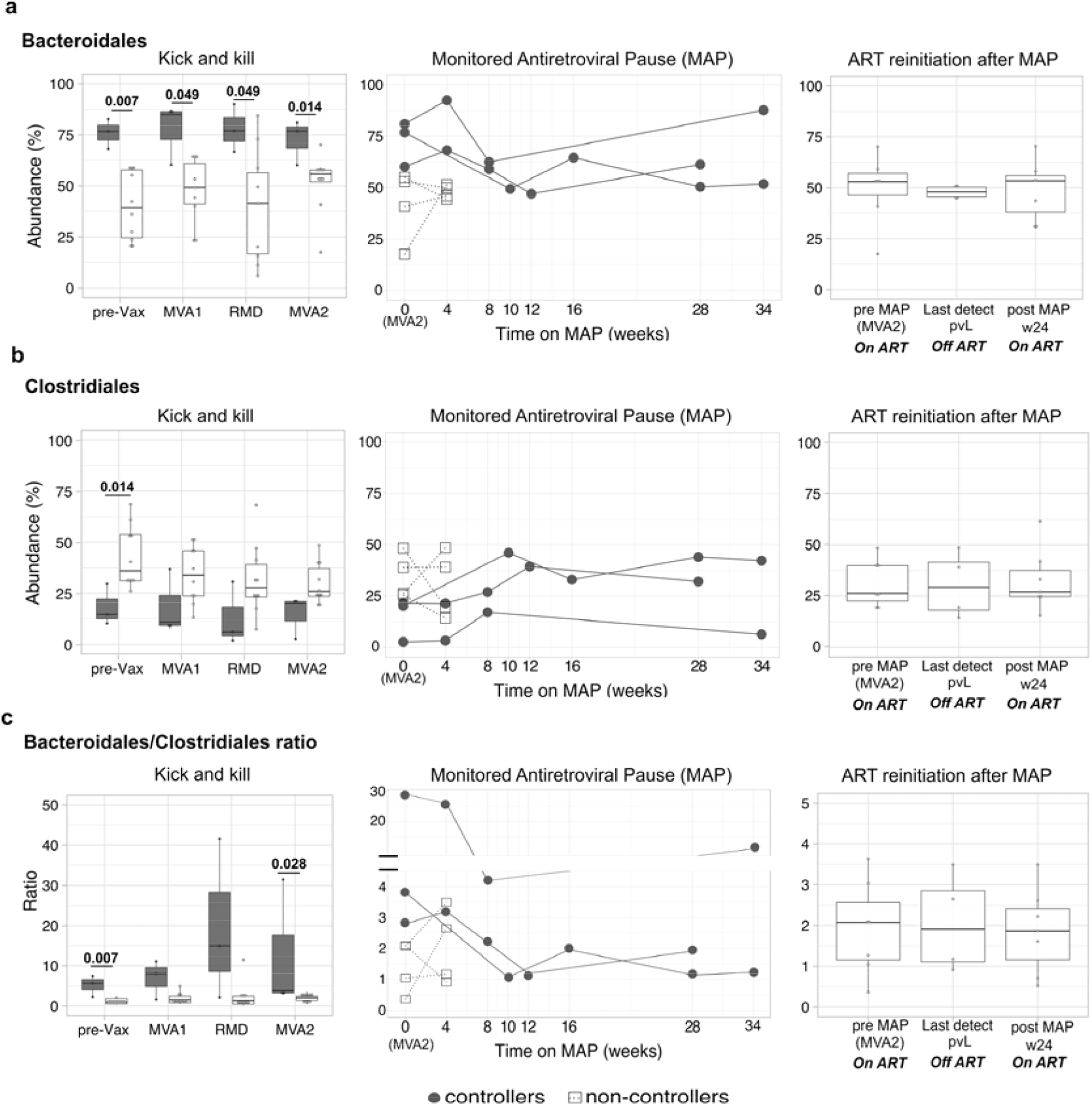
Higher longitudinal *Bacteroidales/Clostridiales* ratio in viremic controllers. Relative abundance expressed as percentage of (**a**), *Bacteroidales*, (**b**) *Clostridiales* and (**c**) their ratio in controllers (gray) and non-controllers (white) are represented by boxplots (left and right vertical panels) and line plots (middle vertical panels). In line plots, values for each subject are illustrated by white squares (non-controllers) and grey dots (viremic controllers). Boxplots show the median (horizontal black line) and interquartile range between the first and third quartiles (25^th^ and 75^th^, respectively). Third vertical panels show non-controllers before ART interruption (pre-MAP, n=7), last timepoint on MAP before ART resumption (Last detect pVL, n=4) and 24 weeks after ART resumption (post MAP w4, n=7). Abbreviations: MAP, monitored antiretroviral pause; pre-Vax, baseline (1 day before first MVA vaccination); MVA1, 1 week after first MVA vaccination; RMD, 1 week after third romidepsin infusion; MVA2, 4 weeks after second MVA vaccination.

### Viremic controllers display lower gut microbial diversity and richness over the intervention

Controllers had lower microbial gene counts than non-controllers at the study entry and throughout the study intervention, although such differences lost statistical significance in the RMD and MVA2 assessments (Fig. 2a). Intra-individual diversity (Shannon index) also remained numerically lower in controllers, but differences were not statistically significant (Fig. 2b). During the MAP, gut microbial diversity increased around weeks 8-10 in controllers, and remained stable thereafter (Figs. 2a-b). No statistically significant differences were found in microbial diversity following ART re-initiation (Figs. 2a-b). Using the Bray-Curtis index, controllers exhibited lower beta-diversity and higher similarity, particularly already at the study entry (Additional Figures: Figure S7), and showed less intra-host longitudinal evolution (Fig. 2c) than non-controllers (PERMANOVA, *r*^2^ = 0.591, *p* = 0.001). Whereas the gut microbiome composition of controllers was significantly different from that of non-controllers (PERMANOVA, *r*^2^ =0.112, *p*=0.001), no significant longitudinal differences were observed within each group (PERMANOVA, *r*^2^ =0.043, *p*=0.815), suggesting that the combined intervention with MVA.HIVconsv vaccines and three weekly low-dose infusion of romidepsin did not significantly alter the gut-microbiome composition (Fig. 2c). Of note, results did not change after removing B07 from the non-controller arm (Additional Figures: Figure S8).

**Fig 2.**
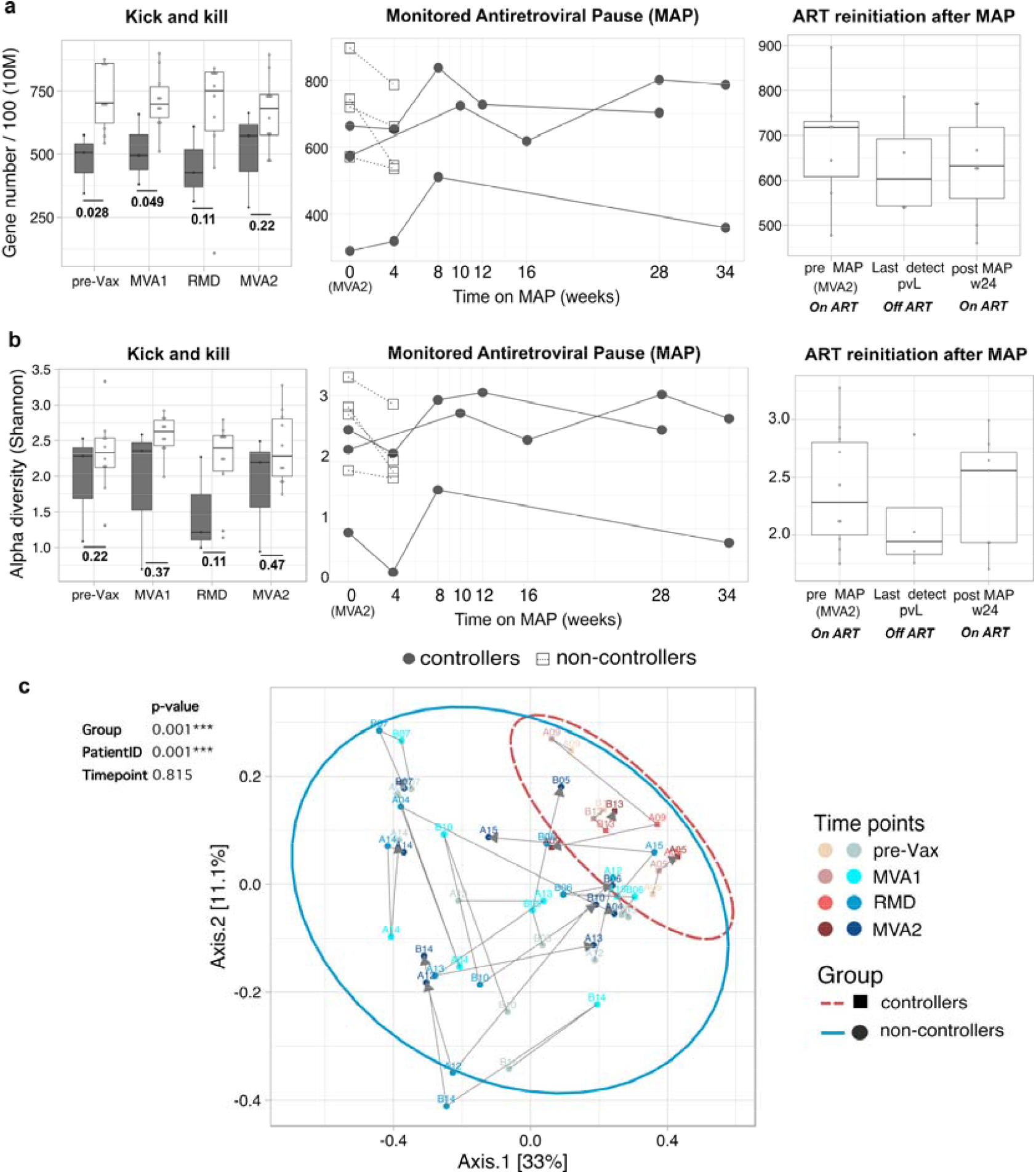
Lower microbial diversity and richness in controllers. Longitudinal (**a**) microbial gene richness at 10 million (10M filtered reads) down-sampling size and (**b**) alpha diversity based on Shannon index in viremic controllers (gray) and non-controllers (white). **c**, Principal coordinates analysis (PCoA) of microbial diversity based on Bray-Curtis distances at pre-vaccination and during ‘kick and kill’ intervention. Proportion of variance explained by each principal coordinate axis is reported in the corresponding axis label. Subjects per each group are represented by squares (controllers) and circles (non-controllers). Each point stands for one subject, color coded by group and time point. The increase in purple (controllers) and blue (non-controllers) colors reflects sequential time points from baseline (pre-Vax) to the second vaccine administration (MVA2). Ellipses delineate the distribution of points per each group. Gray arrows link directional changes in bacterial abundance throughout the kick and kill intervention from baseline (pre-Vax). PERMANOVA statistical analysis of samples grouped by Group, PatientID (patient internal identifier) and time point is shown on the top of the panel. Abbreviations: MAP, monitored antiretroviral pause; pre-Vax, baseline (1 day before first MVA vaccination); MVA1, 1 week after first MVA vaccination; RMD, 1 week after third romidepsin infusion; MVA2, 4 weeks after second MVA vaccination.

### Longitudinal microbial metabolic pathways differ between viremic controllers and non-controllers

Functional profiling based on HUMAnN2[26] identified 28 differential metabolic pathways between controllers and non-controllers at study entry (unadjusted *p* <0.05, Wilcoxon test) (Additional Figures: Figure S9 and S10a). Twelve out of the 28 pathways identified at study entry were differentially abundant throughout the intervention (Additional Figures: Figure S10b). Controllers were enriched in ‘fatty acid and lipid biosynthesis’ and ‘amino acid biosynthesis’ pathways. Conversely, metabolic pathways overrepresented in non-controllers included ‘methanogenesis from H_2_ and CO_2_’, ‘carbohydrate biosynthesis’ and ‘generation of precursor metabolite and energy’. Longitudinal variations of such metabolic pathways during MAP and after ART re-initiation are shown (Additional Figures: Figure S11), although low numbers did not allow for statistical testing. The ‘methanogenesis from H_2_ and CO_2_’ pathway was the most discriminant feature between the two groups (fold-change=11.5, *p*=0.04). Consistently, methanogenic archaea (*Methanobrevibacter smithii* and *Methanosphaera stadtmanae*) were detected in most non-controllers and were rare or absent in controllers (Additional Figures: Figure S12). Taken together, these data show that differences between controllers and non-controllers emerged from resident microbial communities, before any intervention was started in BCN02 study. Thus, subsequent analyses were focused on characterizing further potentially discriminant signatures at study entry.

### Increased *Bacteroidales/Clostridiales* ratio in viremic controllers negatively correlated with longitudinal HIV-1 viral reservoir size

The *Bacteroidales/Clostridiales* ratio inversely and significantly correlated with longitudinal total CD4^+^ T cell-associated HIV-1 DNA measured at study entry (rho□= -0.6, *p*□=□0.03) and over the “kick & kill” intervention, whereas an opposite trend was observed for gene richness (rho= 0.65, *p*□=□0.01 at study entry) (Fig. 3a). Alpha-diversity (Shannon index) exhibited weak positive correlation with the viral reservoir, being the correlation not significant. In line with the gut microbiota patterns found in controllers, the ratio *Bacteroidales/Clostridiales* showed a strong negative correlation with gene richness (r□ho= - 0.87, *p*□=□0.0001) (Fig. 3a). In the longitudinal comparison, controllers tended to displayed lower viral reservoir size, although differences were not statistically significant (Fig. 3b). A similar trend was observed for cell-associated (CA) HIV-1 RNA (Figs. 3c-d), although stronger correlations were found at RMD and MVA2 timepoints with both *Bacteroidales/Clostridiales* ratio (RMD; rho= -0.76, *p*=0.002 and MVA2; rho= - 0.74, *p*=0.003) and gene richness (RMD; rho= 0.72, *p*=0.005 and MVA2; rho= 0.71, *p*=0.006) (Fig. 3c). Moreover, in the longitudinal comparison, controllers displayed significantly lower CA HIV-1 RNA at RMD and MVA2 (*p*=0.03) (Fig. 3d). A set of clinical and vaccine-response variables was screened for association with gut microbial signatures. Absolute CD4^+^ T-cell count before ART initiation was the only factor significantly associated with the *Bacteroidales/Clostridiales* ratio (rho= 0.65, *p*□=□0.01) and gene richness (r□ho= -0.62, *p*□=□0.02), whereas a strong and inverse correlations was found between the Shannon index and CD4/CD8 ratio at BCN02 study entry (rho□= 0.9, *p*□=□2.83e-05) (Additional Figures: Figure S13).

**Fig 3.**
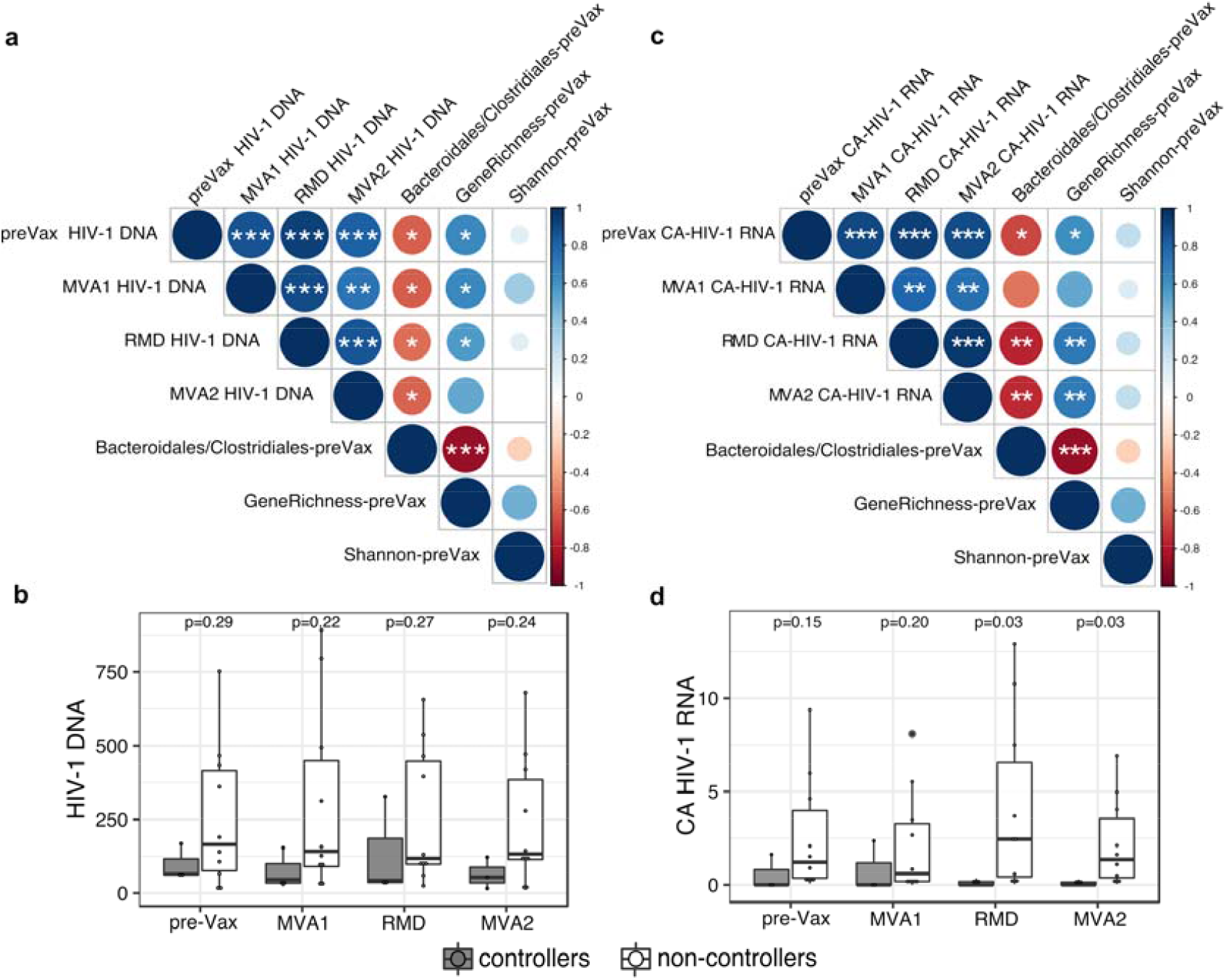
Associations between HIV-1 reservoir size and gut microbial signatures. Spearman’s correlations between gut microbial signatures (ratio *Bacteroidales/Clostridiales*, gene richness and alpha-diversity Shannon index) and longitudinal (**a**) HIV-1 DNA (HIV-1 DNA copies/10^6^ CD4^+^ T-cells) and (**c**) cell-associated (CA) HIV-1 RNA (HIV-1/TBP relative expression). Positive correlations are indicated in blue and negative correlations, in red. Color and size of the circles indicate the magnitude of the correlation. White asterisks indicate significant correlations (\**p* < 0.05; \*\**p* < 0.01; \*\*\**p* < 0.001, Benjamini–Hochberg adjustment for multiple comparisons). Boxplots showing longitudinal comparison of (**b**) HIV-1 DNA and (**d**) cell-associated (CA) HIV-1 RNA between controllers and non-controllers. Abbreviations: MAP, monitored antiretroviral pause; pre-Vax, baseline (1 day before first MVA vaccination); MVA1, 1 week after first MVA vaccination; RMD, 1 week after third romidepsin infusion; MVA2, 4 weeks after second MVA vaccination.

### Distinct bacterial protein signatures associated with viremia control

Baseline metaproteome analysis identified 15,214 bacterial proteins, annotated to 24 unique orders and 69 genera across samples. The abundance of total *Clostridiales* or *Bacteroidales* was not different between groups (Figs. 4a-c). However, several *Clostridiales* genera were decreased in controllers, i.e.: *Eubacterium* (−3.71%; *p*=0.03), *Pseudoflavonifractor* (−0.49%; *p*=0.049), *Oscillibacter* (−0.14%; *p*=0.07), whereas *Blautia* was increased (+5.02%; *p*=0.03) (Fig. 4a). At the genus level, the relative abundance of *Erysipelotrichales* (−0.28%; *p*=0.07), and *Coprobacillus* (−0.22%; *p*=0.07) showed a decreasing trend in controllers, although differences were not statistically significant (Fig. 4b). Unbiased hierarchical clustering showed protein differences (*p*<0.025) between groups (Fig. 4d). Viremic controllers were enriched in bacterial proteins from *Blautia* and *Ruminococcus*, and depleted in proteins derived from other *Clostridiales* such as *Clostridium, Eubacterium, Coprococcus, Faecalibacterium, Oscillibacter, and Pseudoflavinofactor*. Pathways associated with *Blautia* included galactose, starch/sucrose and glyoxylate/dicarboxylate metabolism as well as ribosome activity. (Fig. 4e). Butyrate and other short-chain fatty acid metabolism pathways were similar in both groups (Figs. 4d-e).

**Fig 4.**
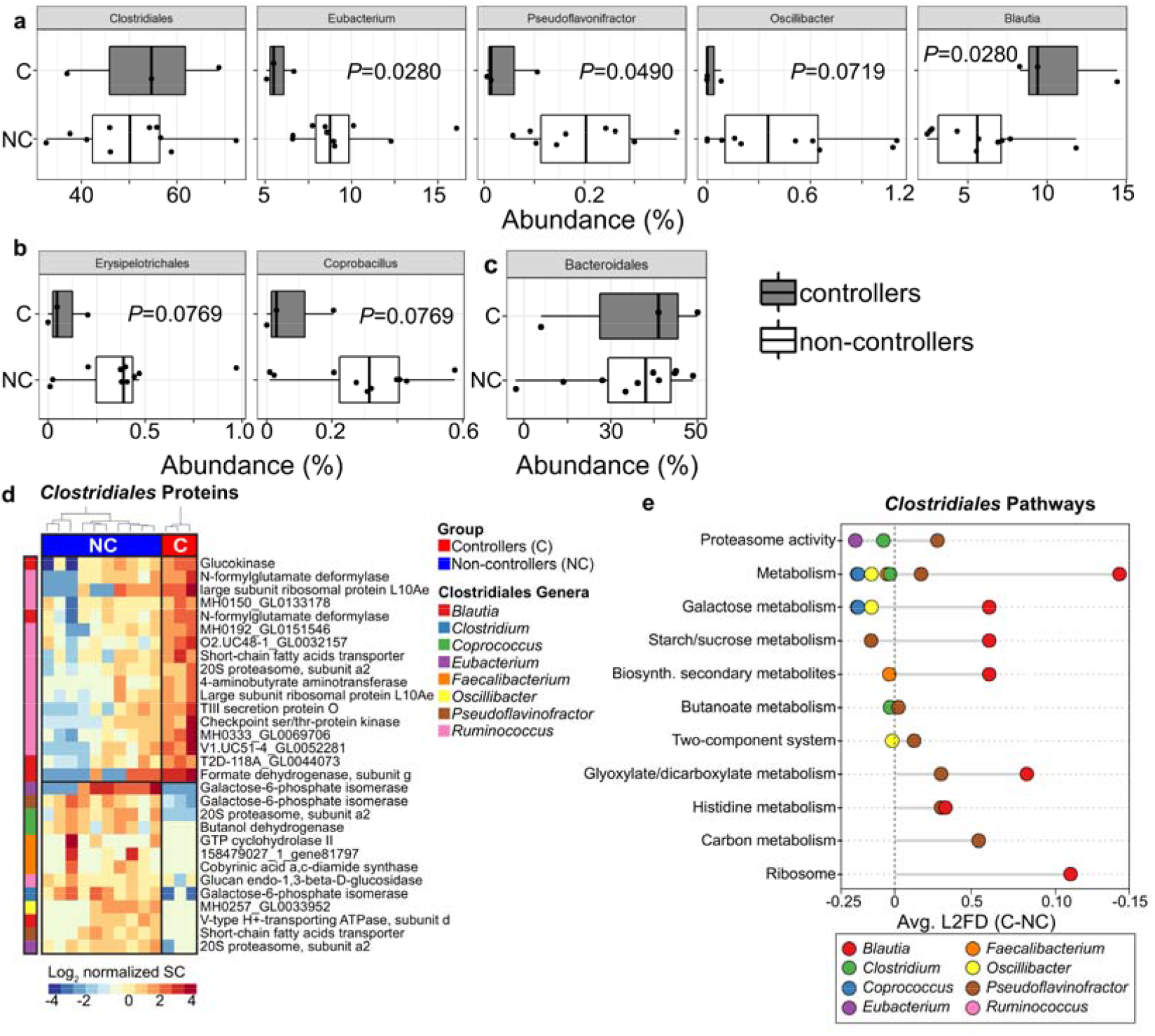
Baseline metaproteomic signatures associated with HIV control after ART interruption. **a-c**, Metaproteomic analysis of gut microbiome in BNC02 participants. Proteins from *Clostridiales* genera and *Erysipelorichales* were relatively underabundant in controllers compared to non-controllers at baseline pre-vaccination. No differences in *Clostridiales* or *Bacteroidales* proteome at the order-level were observed. **d**, Baseline levels of *Clostridiales* proteins (96 proteins) distinguished controllers from non-controllers. Overabundant proteins belonged to the *Blautia* and *Ruminococcus* genera, while under-abundant proteins belonged to the *Clostridium, Coprococcus, Eubacterium, Faecalibacterium, Oscillibacter* and *Pseudoflavinofactor* genera. **e**, Functional annotation of *Clostridiales* bacterial proteins using KEGG gene ontology identified differences in cellular metabolism pathways at baseline between groups. Abbreviations: C=controllers, NC=non-controllers, SC=spectral count.

### Increased baseline immune activation and inflammatory response gene expression in viremic controllers

Full-PBMC gene expression analysis detected a total of 27,426 transcripts at baseline, after filtering for low-expressed genes. Using DESeq2[30], a total of 31 differentially expressed genes (DEGs) were identified (log_2_ FoldChange = 0 and BH-adjusted *p*-value < 0.1), of which 15 and 16 were upregulated in controllers and non-controllers, respectively (Fig. 5a). The proportion of protein-coding genes, pseudogenes, non-coding transcripts, TEC (genes to be experimentally confirmed), and long-intergenic noncoding genes within DEGs was 61.29%, 12.90%, 12.90%, 9.68 and 3.23%, respectively (Additional Table 1). We found 10 and 3 DEGs showing log_2_FC above 2 and 4, respectively; moreover 8 out 10 genes with log_2_FC >2 at a Benjamini-Hochberg-adjusted *p*-value significance < 0.05 (Fig. 5a and Additional Table 1). Hierarchical clustering based on transcriptional DEG profiles showed that controllers grouped together, while non-controllers separated into two distinct expression groups (Additional Fig. 13a). Upregulated genes in non-controllers included 11 transcripts with unknow function (Additional Table 2), which were excluded from downstream analyses. The most statistically differentially-expressed gene was myeloperoxidase (*MPO*, adjusted *p*-value = 1.38e−06), followed by a member of the folate receptor family (*FOLR3*, adjusted *p*-value = 2.68e−05) (Additional Figs. 13b-c and Additional Table 2). Interestingly, both neutrophil-related transcripts *MPO* and *FOLR3* were upregulated in controllers (Fig. 5b) and have been implicated in immune response signaling and regulation of inflammatory processes[33,34]. Other transcripts with known associations to host defense and neutrophil-mediated immunity and higher expression in controllers included defensin alpha 1 and 4 (*DEFA1, DEFA4*), bactericidal permeability-increasing protein (*BPI*), cathepsin G (*CTSG*) and neutrophil elastase (*ELANE*) (Fig. 5b and Additional Table 2). Gene Ontology (GO) enrichment analysis identified a number of biological processes associated with genes highly expressed in controllers (Additional Table 3). To reduce redundancy, the complete list of GO terms was collapsed into representative subclasses using REVIGO. Many of these functions were associated with immune response, such as neutrophil-mediate immunity, leukocyte degranulation and antimicrobial humoral response (Fig. 5c).

**Fig 5.**
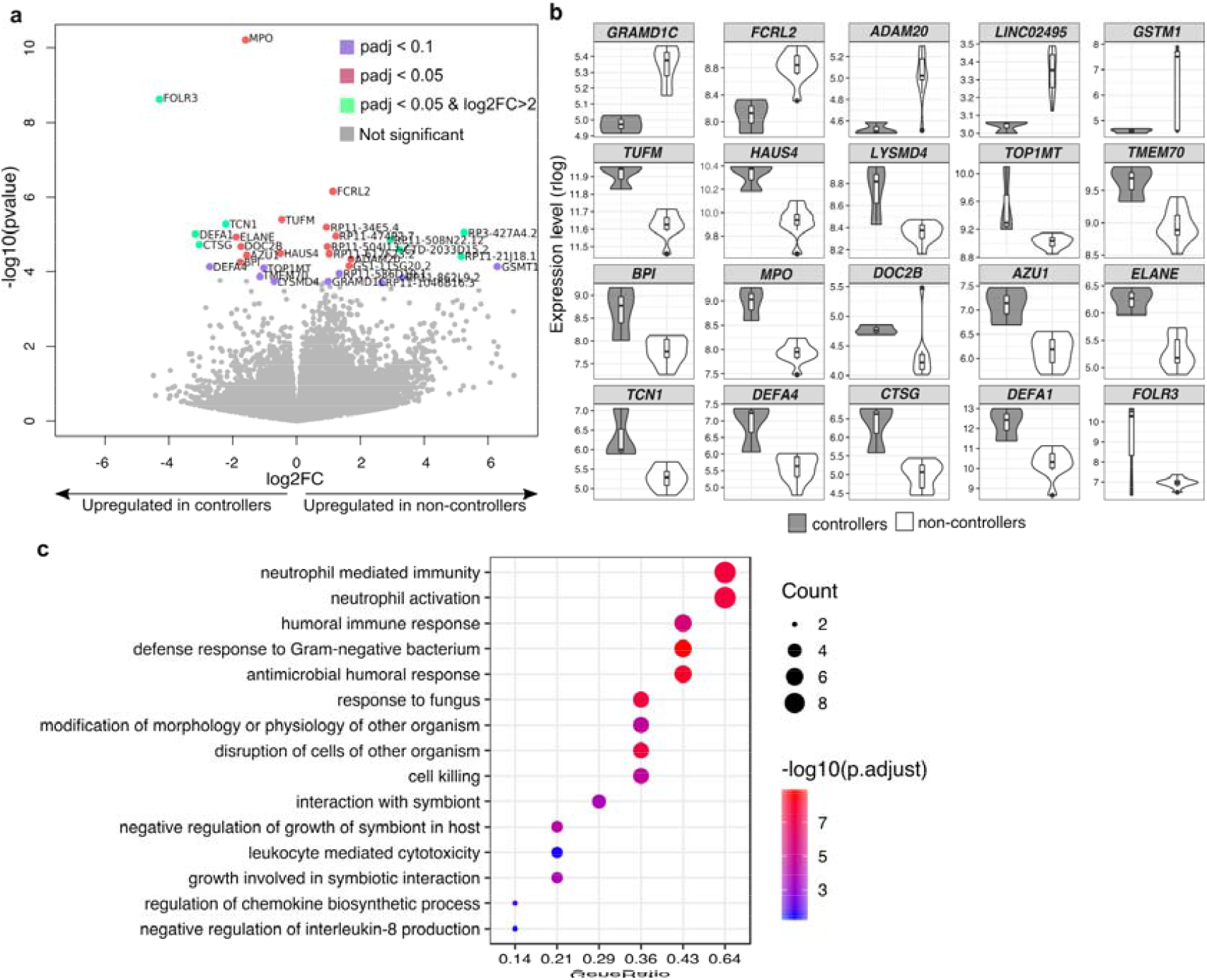
Baseline functional enrichment in levels of immune activation and inflammatory response in viremic controllers. **a**, Volcano plot of differentially expressed genes between controllers and non-controllers at baseline (pre-Vax) with adjusted *p-*value < 0.1 (violet dots), adjusted *p-*value < 0.05 (red dots) and log2 (FoldChange) > 2 and adjusted *p-*value < 0.05 (green dots). Gray-colored dots represent genes not displaying statistical significance (adjusted *p*-value > 0.1). The log2 Fold Change on the x-axis indicates the magnitude of change, while the −log10 (p-value) on the y-axis indicates the significance of each gene. A total of 31 genes showed significant differential expression (adjusted *p*-value < 0.1): 15 and 16 upregulated in controllers and non-controllers, respectively. **b**, Violin plots showing relative expression levels (rlog, regularized log transformation) of differentially expressed genes with functional annotation. **c**, Gene ontology (GO) enrichment analysis of upregulated genes in Controllers. In the y-axis, only representative enriched GO terms (biological process) are reported (terms obtained after redundancy reduction by REVIGO). X-axis reports the percentage of genes in a given GO terms, expressed as ‘Gene ratio’. The color key from blue to red indicates low to high Bonferroni-adjusted log 10 *p*-value. Dot sizes are based on the “count” (genes) associated to each GO term. Significantly enriched GO terms, number of genes associated to each GO term and adjusted *p*-values are provided in Additional Table 3.

### Increased baseline inflammation-related plasma proteins in viremic controllers

Soluble factors in plasma from the 92 inflammation-related protein panel were assessed using the Proximity Extension Assay (see Methods in Additional Text). Plasma protein levels did not independently separate controllers from non-controllers using unbiased hierarchical clustering or principal component analysis (Additional Figs. 14a-b). Of the 92 plasma proteins characterized, only 7 were differentially expressed (Wilcoxon, uncorrected *p*<0.05) and were increased in controllers (Additional Fig. 14c): adenosine deaminase ADA (p=0.012), decoy receptor osteoprotegerin OPG (p=0.024), self-ligand receptor of the signaling lymphocytic activation molecule family SLAMF-1 (p = 0.048), chemokines CCL23, CCL28, MCP-2 (p = 0.048) and the neurotrophic factor NT-3 (p = 0.048) (Additional Fig. 14d).

### Integration analysis between *Bacteroidales/Clostridiales* ratio, host immune activation-related transcripts, bacterial proteins and HIV-1 reservoir size

The *Bacteroidales / Clostridiales* ratio was positively correlated (Spearman rho=0.55 and *q*-value < 0.05) with DEGs involved in inflammatory response and immune system activation, including *DEFA1, DEFA4, TOP1MT, CTSG, MPO, AZU1, ELANE* (Fig. 6a), Spearman rho and *q* value are given in Additional Dataset 1. Functional enrichment analysis including all transcripts significantly correlated with the ratio *Bacteroidales/Clostridiales* (n. transcripts=453, *q*-value <0.05, see Additional Dataset 1) identified four main functional clusters related to neutrophil activation, disruption of cells of other organism, antimicrobial humoral response and regulation of immune response to tumor cell (Additional Fig. 15a and Additional Table 4). Of these, the *Bacteroidales / Clostridiales* ratio strongly correlated (rho >0.8) with transcripts within such functional clusters, which were highly expressed in controllers (Fig. 6b).

**Fig 6.**
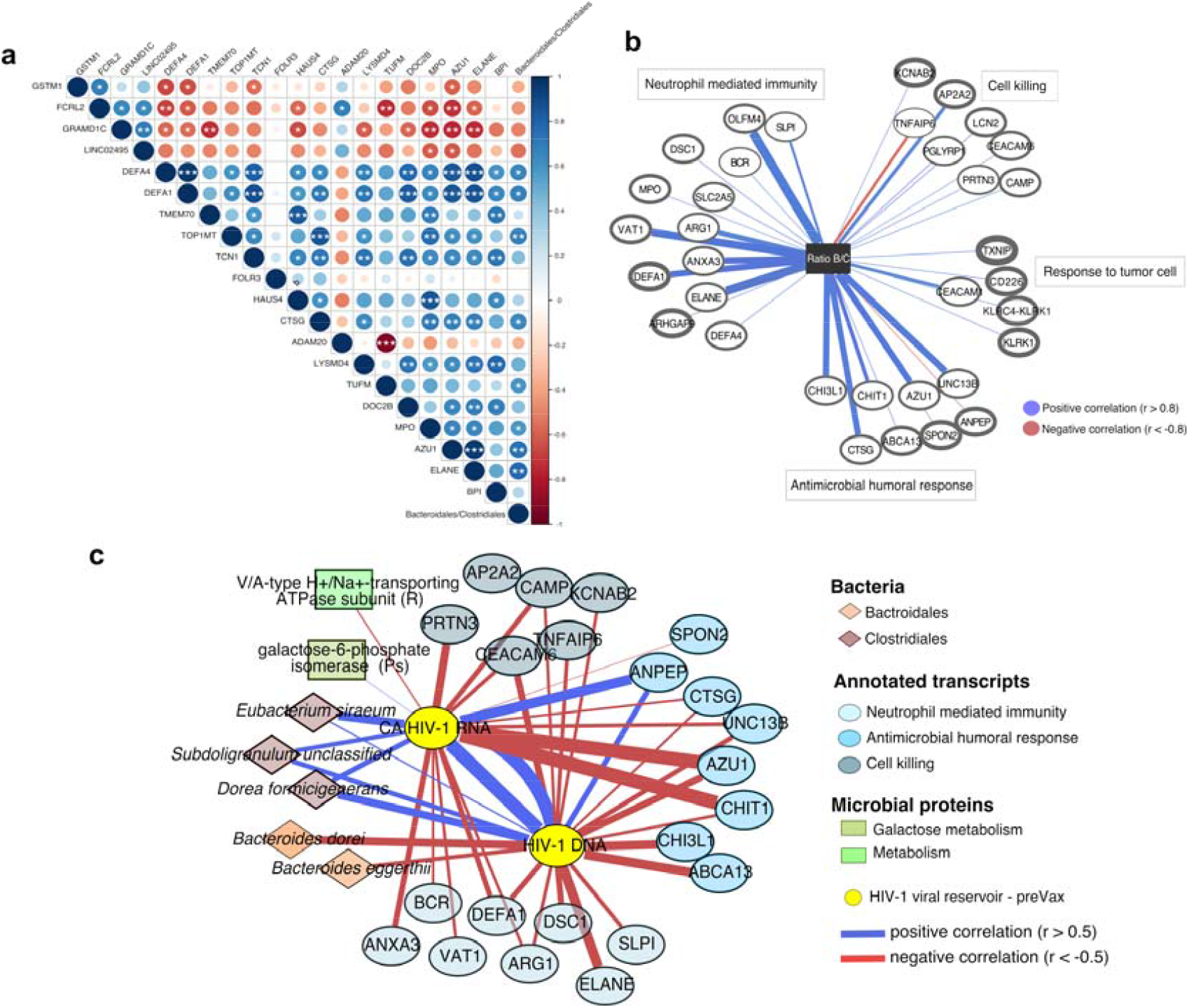
Integrative analysis of baseline gut microbial signatures, immune activation-related transcripts, bacterial proteins and HIV-1 reservoir. **a**, Spearman’s correlations between the ratio *Bacteroidales/Clostridiales* and DEGs (annotated transcripts). Positive correlations are indicated in blue and negative correlations, in red. Color and size of the circles indicate the magnitude of the correlation. White asterisks indicate significant correlations (*p < 0.05; **p < 0.01; ***p < 0.001, Benjamini–Hochberg adjustment for multiple comparisons). **b**, Network visualizing significant Spearman’s correlations between the ratio *Bacteroidales/Clostridiales* and transcripts involved in the enrichment analysis described in Additional Table 4. Transcripts are represented as vertices and border width is proportional to transcript expression (log2 |cpmTMM_w0 +1|) in controllers. Edge width indicates the magnitude of correlation. Red and blue edges represent positive and negative correlation, respectively. **c**, Network showing Spearman’s correlation between viral reservoir (CA HIV-1 RNA and HIV-1 DNA), bacterial species within *Bacteroidales* and *Clostrdiales*, human transcripts correlated with the ratio *Bacteroidales / Clostrdiales* and differentially abundant bacterial proteins (*p* ≤ 0.025). Features are showed as vertices and colored by ‘omic’ dataset. Positive and negative correlations are presented as blue and red edges, respectively. Edge width indicates the magnitude of correlation coefficient. Protein-associated bacterial genera are reported in parentheses. Abbreviations: DEGs, differentially expressed genes between controllers and non-controllers; R, *Ruminococcus*; Ps, *Pseudoflavonifactor* and pre-Vax, baseline timepoint (1 day before first MVA vaccination).

Moreover, 61 and 70 out 453 transcripts (Additional Dataset 2) correlated (rho = 0.5) with the baseline CA HIV-1 RNA and HIV-1 DNA, respectively, and were enriched in immune-mediated response functions (Additional Figs. 15b-c and Additional Table 5). In the integrated analysis of metagenomic, transcriptomic and metaproteomic data to identify signatures discriminating between controllers and non-controllers, *Bacteroidales* and *Clostridiales* were clearly separated through the component (Additional Fig. 16a). While *Bacteroidales* clustered with immune activation-related transcripts (*MPO, AZU1, ELANE, TCN1, DEFA1, BPI, DEF4*) and proteins from *Ruminococcus, Blautia, Prevotella* and *Faecalibacterium* genera, *Clostridiales* inversely correlated with these features (Additional Figs. 16a-b). Such associations were assessed at a lower taxonomic scale and confirmed that *Bacteroidales* species (*B. dorei* and *B. eggerthii*) inversely correlated with HIV-1 DNA levels, whereas members of *Clostridiales* (*S. unclassified, D. formicigenerans* and *E. siraeum*) positively correlated with HIV-1 DNA and CA HIV-1 RNA levels. The viral reservoir (HIV-1 DNA and CA HIV-1 RNA) negatively correlated with genes involved in ‘neutrophil mediated immunity’, ‘antimicrobial humoral response’ and ‘cell killing’. Weak correlations were observed with bacterial proteins involved in the regulation of metabolism (Fig. 6c and Additional Dataset 2). Overall, these findings further supported positive associations between *Bacteroidales* abundance and transcripts related to immune response in controllers, which in turn negatively correlated with the viral reservoir size.

## Discussion

In this proof-of-concept study, a longitudinal multi-’omics’ analysis identified the *Bacteroidales/Clostridiales* ratio as a novel gut microbiome signature associated with HIV-1 reservoir size and viremic control during a monitored ART pause. Individuals with high *Bacteroidales/Clostridiales* ratio showed gene expression signatures related to immune activation, particularly neutrophil-mediated immunity and antimicrobial humoral response, which negatively correlated with the viral reservoir size. Our findings largely arise from unsupervised analyses where many other signatures could have emerged, especially given the relatively low number of individuals analyzed. However, they are internally coherent and consistent with a theoretical framework where increased inflammation might contribute to immune-mediated HIV-1 control. They also suggest a putative biomarker for safer ART interruptions in HIV cure studies.

The baseline gut microbiome of controllers was enriched in pro-inflammatory species, such as *P. copri*[35], and depleted in bacteria, traditionally associated with the maintenance of gut homeostasis through production of SCFAs[36], including *R. intestinalis* and *Subdoligranulum spp*. Lower microbial diversity and gene richness in controllers were consistent with a previous work from our group in people living with HIV[37], as well as other studies[38], in which higher gene richness associated with increased levels of butyrate-producing bacteria and methanogenic archaea. Microbial functional enrichment in ‘lipid and fatty acid biosynthesis’ in controllers might be reflective of mechanisms of lipopolysaccharide biosynthesis and production of inflammatory mediators[39] mediated by members of *Bacteroidales*[40]. No discernible longitudinal variations were observed in the gut microbiome of BCN02 participants, in line with a previous evidence in oral typhoid immunization strategy[16]. Of note, the gut microbiome in healthy population has been described generally resilient to perturbations[41]. Taken together, these observations would suggest a trend toward the maintenance of a relative stability in the gut microbiome composition, with resident microbial communities potentially associated with viral control during ART interruption. Baseline metaproteome analysis confirmed that functional differences between controllers and non-controllers were mainly driven by *Clostridiales*, which were actively producing microbial proteins in both groups albeit in distinct functional contexts. Further discriminating baseline signatures linked to increased immune system activation and inflammatory response in controllers emerged from PBMC transcriptome and inflammation-related plasma proteins profiling. It also emerged that the ratio *Bacteroidales/Clostridiales* inversely correlated with the viral reservoir size in terms of HIV-1 DNA and CA HIV-1 RNA. Although controllers did not display significantly lower viral reservoir size compared to non-controllers, these associations are consistent with previous studies suggesting a role of low viral reservoir on ART interruption outcomes[9].

Taken together, these findings suggest that baseline immune activation potentially associated with a microbial shift toward pro-inflammatory bacteria and lower viral reservoir may contribute to sustained post ART interruption HIV-1 control. While there is evidence suggesting a strong impact of the gut microbiota composition on host immune system and inflammatory status[42], the mechanistic basis of how microbial communities may interact with the viral reservoir and, in turn, exert immunomodulatory effects on HIV-1 control during ART interruption remains to be delineated. We speculate that a pre-existing, altered balance of ‘beneficial’ gut microbial groups, such as *Clostridiales*, and concomitant overabundance of pro-inflammatory bacteria would boost host immune system activation, thus triggering a prompt control of rebounding virus, as observed in controllers. In support of this hypothesis, increased abundance of members from *Clostridiales* were previously associated with neutrophilia and lower poliovirus and tetanus-hepatitis B vaccine response[18]. Moreover, baseline transcriptional pro-inflammatory and immune activation signatures were suggested as potential predictors of increased influenza[43], systemic lupus erythematosus[44] and hepatitis B[45] vaccine-induced immune response, with weaker responses in elderly[43,45]. It is thus reasonable to postulate that immune activation prior to vaccination together with microbiome-associated factors may affect vaccine outcomes.

This study has several limitations. Due to eligibility criteria in the parental BCN02 study, there was a limited sample size, and we were unable to include a control arm without the intervention. Therefore, our considerations were narrowed to three individuals that showed viremic control during ART interruption. Bearing these limitations in mind, our results should be interpreted with caution, emphasizing the need of independent validation in randomized and placebo-controlled trials to assess potentially unmeasured confounders and provide further perspectives on factors that might induce gut microbial shifts. Upcoming analyses in larger longitudinal trials, including the recently reported AELIX-002 trial[46], where fecal samples have been stored longitudinally, are expected to validate our findings. These preliminary findings might have important implications in the design of HIV-1 cure intervention trials that include ART interruption. As proposed for other therapeutic areas[47], microbiome-associated predictive patterns could help to optimize patient stratification, thus resulting in more targeted studies and higher efficacy of HIV-1 interventions. In addition, if a given resident microbial community is to be defined that is indeed predictive of viral control during ART interruption, then modulating participants’ gut microbiota before immunization might potentially modulate vaccine responsiveness and ultimately, clinical outcomes. While host-genetics and other vaccine-associated factors as baseline predictors are less amendable, the gut microbiome is potentially modifiable and even transferrable to another host. Strategies manipulating the gut microbiota composition and relative by-products via prebiotics and/or probiotics administration[48] or microbiota engraftment following fecal microbiota transplantation[49] are under intense evaluation[50], albeit with several limitations.

## Conclusions

In conclusion, in this exploratory study, we identified pre-existing gut microbial and immune activation signatures as potential predictors of sustained HIV-1 control in the absence of ART, providing a potential target for future treatment strategies and opening up new avenues for a functional HIV cure.

## Availability of data and materials

Datasets supporting the conclusions of this study are available as Additional information (Additional Datasets). Metagenome and RNA-seq data have been deposited in the European Nucleotide Archive (ENA) and are accessible through ENA accession numbers PRJEB42384 and PRJEB43195. The code and databases used for data analysis are available as Additional information (Additional Code) and at 10.5281/zenodo.4876340.

## Abbreviations

ART: antiretroviral therapy
ATI: antiretroviral therapy interruption
HUMAnN2: HMP unified metabolic analysis network v.2
MAP: monitored antiretroviral therapy pause
MVA: MVA.HIVconsv
RMD: romidepsin
SCFA: short chain fatty acids

## Acknowledgements

The authors thank all volunteers for participating in this study and the BCN02 study group.

## BCN02 Study Group

IrsiCaixa AIDS Research Institute-HIVACAT Hospital Universitari Germans Trias i Pujol, Badalona, Spain: Susana Benet, Christian Brander, Samandhy Cedeño, Bonaventura Clotet, Pep Coll, Anuska Llano, Javier Martinez-Picado, Marta Marszalek, Sara Morón-López, Beatriz Mothe, Roger Paredes, Maria C. Puertas, Miriam Rosás-Umbert, Marta Ruiz-Riol. Fundació Lluita contra la Sida, Infectious Diseases Department, Hospital Universitari Germans Trias i Pujol, Badalona, Spain: Roser Escrig, Silvia Gel, Miriam López, Cristina Miranda, José Moltó, Jose Muñoz, Nuria Perez-Alvarez, Jordi Puig, Boris Revollo, Jessica Toro. Germans Trias i Pujol Research Institute, Badalona, Spain: Ana María Barriocanal, Cristina Perez-Reche. Clinical Pharmacology Unit, Hospital Universitari Germans Trias i Pujol, Badalona, Spain: Magí Farré. Pharmacokinetic/pharmacodynamic modeling and simultation, Institut de Recerca de l’Hospital de la Santa Creu i Sant Pau-IIB Sant Pau, Barcelona, Spain: Marta Valle. Hospital Clinic-HIVACAT, IDIBAPS, University of Barcelona, Barcelona, Spain: Christian Manzardo, Juan Ambrosioni, Irene Ruiz, Cristina Rovira, Carmen Hurtado, Carmen Ligero, Emma Fernández, Sonsoles Sánchez-Palomino, and Jose M. Miró. Projecte dels NOMS-Hispanosida, BCN Checkpoint, Barcelona, Spain: Antonio Carrillo, Michael Meulbroek, Ferran Pujol and Jorge Saz. The Jenner Institute, The Nuffield Department of Medicine, University of Oxford, UK: Nicola Borthwick, Alison Crook, Edmund G. Wee and Tomás□ Hanke.

## Author Contributions

R.P, B.M, J.M, B.C, J.M.M and C.B, conceived and designed the study. B.M, R.P, C.M, J.M.M and C.B recruited the study participants and performed their clinical evaluations. M.P and M.C performed fecal DNA extraction, library preparation and sequencing, under the supervision of M.N.J, Y.G and R.P. B.O and C.D performed PBMC transcriptomics and soluble factors determinations, under the supervision of M.R.R and C.B. M.C.P performed the viral reservoir size determinations, under the supervision of J.M.P. L.N.R, M.D.L, S.K and K.B performed fecal metaproteomics experiments and data analysis, under the supervision of A.Bu. A.Bo performed bioinformatics and statistical analyses of metagenome, transcriptome, soluble factors, clinical and integration data, under the supervision of M.N.J and R.P. L.N.R performed the bioinformatics and statistical analyses of fecal metaproteome data, under the supervision of A.Bu. F.C.M, M.N.J, A.Bo, B.O and Y.G contributed to data management. A.Bo and R.P wrote the paper, which was reviewed, edited and approved by all authors.

## References

1. Volberding PA, Deeks SG. Antiretroviral therapy and management of HIV infection. Lancet. 2010. p. 49–62.

2. Finzi D, Hermankova M, Pierson T, Carruth LM, Buck C, Chaisson RE, et al. Identification of a reservoir for HIV-1 in patients on highly active antiretroviral therapy. Science (80-). 1997;278:1295–300.

3. Siliciano JD, Kajdas J, Finzi D, Quinn TC, Chadwick K, Margolick JB, et al. Long-term follow-up studies confirm the stability of the latent reservoir for HIV-1 in resting CD4+ T cells. Nat Med. 2003;9:727–8.

4. Chun TW, Davey RT, Engel D, Lane HC, Fauci AS. AIDS: Re-emergence of HIV after stopping therapy. Nature. 1999;401:874–5.

5. Namazi G, Fajnzylber JM, Aga E, Bosch RJ, Acosta EP, Sharaf R, et al. The control of HIV after antiretroviral medication pause (CHAMP) study: Posttreatment controllers identified from 14 clinical studies. J Infect Dis. 2018;218:1954–63.

6. Turnbull EL, Wong M, Wang S, Wei X, Jones NA, Conrod KE, et al. Kinetics of Expansion of Epitope-Specific T Cell Responses during Primary HIV-1 Infection. J Immunol. 2009;182:7131–45.

7. Liu MKP, Hawkins N, Ritchie AJ, Ganusov V V., Whale V, Brackenridge S, et al. Vertical T cell immunodominance and epitope entropy determine HIV-1 escape. J Clin Invest. 2013;123:380–93.

8. Posteraro B, Pastorino R, Di Giannantonio P, Ianuale C, Amore R, Ricciardi W, et al. The link between genetic variation and variability in vaccine responses: Systematic review and meta-analyses. Vaccine. 2014;32:1661–9.

9. Li JZ, Etemad B, Ahmed H, Aga E, Bosch RJ, Mellors JW, et al. The size of the expressed HIV reservoir predicts timing of viral rebound after treatment interruption. AIDS. 2016;30:343–53.

10. Zimmermann P, Curtis N. The influence of the intestinal microbiome on vaccine responses. Vaccine. 2018. p. 4433–9.

11. Ciabattini A, Olivieri R, Lazzeri E, Medaglini D. Role of the microbiota in the modulation of vaccine immune responses. Front. Microbiol. 2019.

12. Sui Y, Lewis GK, Wang Y, Berckmueller K, Frey B, Dzutsev A, et al. Mucosal vaccine efficacy against intrarectal SHIV is independent of anti-Env antibody response. J Clin Invest. 2019;129:1314–28.

13. Pantaleo G, Janes H, Karuna S, Grant S, Ouedraogo GL, Allen M, et al. Safety and immunogenicity of a multivalent HIV vaccine comprising envelope protein with either DNA or NYVAC vectors (HVTN 096): a phase 1b, double-blind, placebo-controlled trial. Lancet HIV. 2019;6:e737–49.

14. Pantaleo G, Janes H, Tomaras G, Montefiori D, Frahm N, Grant S, et al. Comparing different priming strategies to optimize HIV vaccine antibody responses: results from HVTN 096/EV04 (NCT01799954). AIDS Res Hum retroviruses Conf 2nd HIV Res Prev Conf HIVR4P 2016 United states [Internet]. 2016;32:68. Available from: https://www.cochranelibrary.com/central/doi/10.1002/central/CN-01646903/full

15. Cram JA, Fiore-Gartland AJ, Srinivasan S, Karuna S, Pantaleo G, Tomaras GD, et al. Human gut microbiota is associated with HIV-reactive immunoglobulin at baseline and following HIV vaccination. PLoS One. 2019;14.

16. Eloe-Fadrosh EA, McArthur MA, Seekatz AM, Drabek EF, Rasko DA, Sztein MB, et al. Impact of Oral Typhoid Vaccination on the Human Gut Microbiota and Correlations with S. Typhi-Specific Immunological Responses. PLoS One. 2013;8.

17. Harris VC, Armah G, Fuentes S, Korpela KE, Parashar U, Victor JC, et al. Significant Correlation Between the Infant Gut Microbiome and Rotavirus Vaccine Response in Rural Ghana. J Infect Dis. 2017;215:34–41.

18. Huda MN, Lewis Z, Kalanetra KM, Rashid M, Ahmad SM, Raqib R, et al. Stool microbiota and vaccine responses of infants. Pediatrics. 2014;134.

19. Julg B, Dee L, Ananworanich J, Barouch DH, Bar K, Caskey M, et al. Recommendations for analytical antiretroviral treatment interruptions in HIV research trials—report of a consensus meeting. Lancet HIV. 2019. p. e259–68.

20. Mothe B, Rosás-Umbert M, Coll P, Manzardo C, Puertas MC, Morón-López S, et al. HIVconsv Vaccines and Romidepsin in Early-Treated HIV-1-Infected Individuals: Safety, Immunogenicity and Effect on the Viral Reservoir (Study BCN02). Front Immunol. 2020;11.

21. Wei DG, Chiang V, Fyne E, Balakrishnan M, Barnes T, Graupe M, et al. Histone Deacetylase Inhibitor Romidepsin Induces HIV Expression in CD4 T Cells from Patients on Suppressive Antiretroviral Therapy at Concentrations Achieved by Clinical Dosing. PLoS Pathog. 2014;10.

22. Mothe B, Manzardo C, Sanchez-Bernabeu A, Coll P, Morón-López S, Puertas MC, et al. Therapeutic Vaccination Refocuses T-cell Responses Towards Conserved Regions of HIV-1 in Early Treated Individuals (BCN 01 study). EClinicalMedicine. 2019;11:65–80.

23. Létourneau S, Im EJ, Mashishi T, Brereton C, Bridgeman A, Yang H, et al. Design and pre-clinical evaluation of a universal HIV-1 vaccine. PLoS One. 2007;2.

24. Truong DT, Franzosa EA, Tickle TL, Scholz M, Weingart G, Pasolli E, et al. MetaPhlAn2 for enhanced metagenomic taxonomic profiling. Nat. Methods. 2015. p. 902–3.

25. Li J, Wang J, Jia H, Cai X, Zhong H, Feng Q, et al. An integrated catalog of reference genes in the human gut microbiome. Nat Biotechnol. 2014;32:834–41.

26. Franzosa EA, McIver LJ, Rahnavard G, Thompson LR, Schirmer M, Weingart G, et al. Species-level functional profiling of metagenomes and metatranscriptomes. Nat Methods. 2018;15:962–8.

27. Klatt NR, Cheu R, Birse K, Zevin AS, Perner M, Noël-Romas L, et al. Vaginal bacteria modify HIV tenofovir microbicide efficacy in African women. Science (80-). 2017;356:938–45.

28. Dobin A, Davis CA, Schlesinger F, Drenkow J, Zaleski C, Jha S, et al. STAR: ultrafast universal RNA-seq aligner. Bioinformatics [Internet]. 2013 [cited 2017 Feb 22];29:15–21. Available from: http://www.ncbi.nlm.nih.gov/pubmed/23104886

29. Li B, Dewey CN. RSEM: Accurate transcript quantification from RNA-Seq data with or without a reference genome. BMC Bioinformatics. 2011;12.

30. Love MI, Huber W, Anders S. Moderated estimation of fold change and dispersion for RNA-seq data with DESeq2. Genome Biol. 2014;15.

31. Assarsson E, Lundberg M, Holmquist G, Björkesten J, Thorsen SB, Ekman D, et al. Homogenous 96-plex PEA immunoassay exhibiting high sensitivity, specificity, and excellent scalability. PLoS One. 2014;9.

32. Rohart F, Gautier B, Singh A, Lê Cao KA. mixOmics: An R package for ‘omics feature selection and multiple data integration. PLoS Comput Biol. 2017;13.

33. Aratani Y. Myeloperoxidase: Its role for host defense, inflammation, and neutrophil function. Arch. Biochem. Biophys. 2018. p. 47–52.

34. Holm J, Hansen SI. Characterization of soluble folate receptors (folate binding proteins) in humans. Biological roles and clinical potentials in infection and malignancy. Biochim. Biophys. Acta - Proteins Proteomics. 2020.

35. Iljazovic A, Roy U, Gálvez EJC, Lesker TR, Zhao B, Gronow A, et al. Perturbation of the gut microbiome by Prevotella spp. enhances host susceptibility to mucosal inflammation. Mucosal Immunol. 2020;

36. Lopetuso LR, Scaldaferri F, Petito V, Gasbarrini A. Commensal Clostridia: Leading players in the maintenance of gut homeostasis. Gut Pathog. 2013.

37. Guillén Y, Noguera-Julian M, Rivera J, Casadellà M, Zevin AS, Rocafort M, et al. Low nadir CD4+ T-cell counts predict gut dysbiosis in HIV-1 infection. Mucosal Immunol. 2019;12:232–46.

38. Le Chatelier E, Nielsen T, Qin J, Prifti E, Hildebrand F, Falony G, et al. Richness of human gut microbiome correlates with metabolic markers. Nature. 2013;500:541–6.

39. Medzhitov R, Janeway C. J. Advances in immunology: Innate immunity. N Engl J Med. 2000;343:338–44.

40. d’Hennezel E, Abubucker S, Murphy LO, Cullen TW. Total Lipopolysaccharide from the Human Gut Microbiome Silences Toll-Like Receptor Signaling. mSystems. 2017;2.

41. Fassarella M, Blaak EE, Penders J, Nauta A, Smidt H, Zoetendal EG. Gut microbiome stability and resilience: Elucidating the response to perturbations in order to modulate gut health. Gut. 2020.

42. Zheng D, Liwinski T, Elinav E. Interaction between microbiota and immunity in health and disease. Cell Res. 2020. p. 492–506.

43. Avey S, Cheung F, Fermin D, Frelinger J, Gaujoux R, Gottardo R, et al. Multicohort analysis reveals baseline transcriptional predictors of influenza vaccination responses. Sci Immunol. 2017;2.

44. Kotliarov Y, Sparks R, Martins AJ, Mulè MP, Lu Y, Goswami M, et al. Broad immune activation underlies shared set point signatures for vaccine responsiveness in healthy individuals and disease activity in patients with lupus. Nat Med. 2020;26:618–29.

45. Fourati S, Cristescu R, Loboda A, Talla A, Filali A, Railkar R, et al. Pre-vaccination inflammation and B-cell signalling predict age-related hyporesponse to hepatitis B vaccination. Nat Commun. 2016;7.

46. Lucia Bailon, Anuska Llano, Samandhy Cedeño, Miriam B. Lopez, Yovaninna Alarcón-Soto, Pep Coll, Àngel Rivero, Anne R. Leselbaum, Ian McGowan, Devi SenGupta, Bonaventura Clotet, Christian Brander, Jose Molt BM. A placebo-controlled ATI trial of HTI vaccines in early treated HIV infection. CROI - Virtual Conf Retroviruses Opportunistic Infect. 2021.

47. Boessen R, Heerspink HJL, De Zeeuw D, Grobbee DE, Groenwold RHH, Roes KCB. Improving clinical trial efficiency by biomarker-guided patient selection. Trials. 2014;15.

48. Wilson NL, Moneyham LD, Alexandrov AW. A Systematic Review of Probiotics as a Potential Intervention to Restore Gut Health in HIV Infection. J Assoc Nurses AIDS Care. 2013;24:98–111.

49. Vujkovic-Cvijin I, Rutishauser RL, Pao M, Hunt PW, Lynch S V., McCune JM, et al. Limited engraftment of donor microbiome via one-time fecal microbial transplantation in treated HIV-infected individuals. Gut Microbes. 2017;8:440–50.

50. Rosel-Pech C, Chávez-Torres M, Bekker-Méndez VC, Pinto-Cardoso S. Therapeutic avenues for restoring the gut microbiome in HIV infection. Curr. Opin. Pharmacol. 2020. p. 188–201.

