## Additional for "Gut Microbiome Signatures Linked to HIV-1 Reservoir Size and Viremia Control": Additional Figures.docx

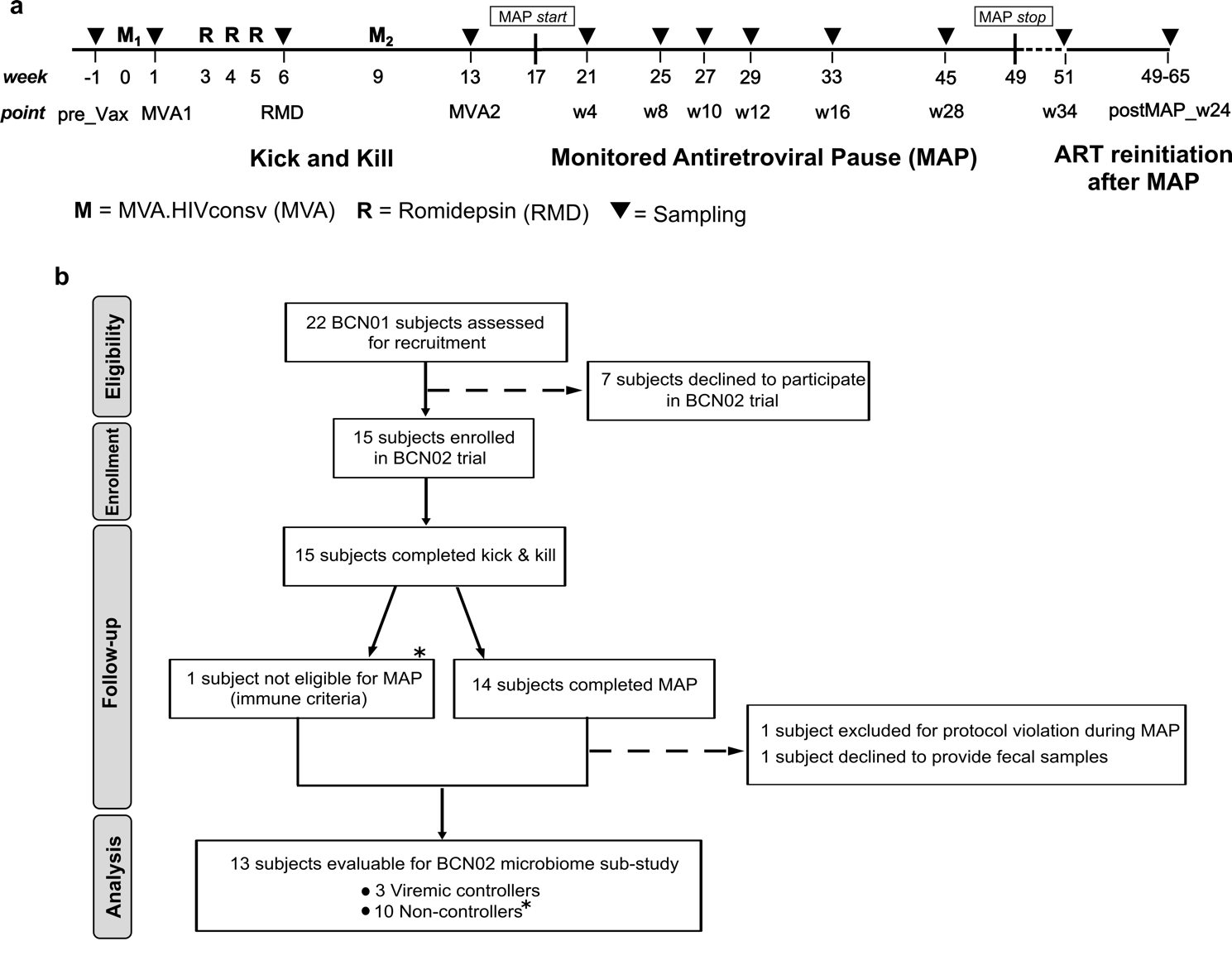

**Figure S1**. **BCN02-microbiome study design and sample collection strategy. a**, BCN02 trial participants were immunized with two doses of MVA.HIVconsv vaccine, before (M_1_) and after (M_2_) three weekly-doses of romidepsin (RMD) followed by a monitored antiretroviral pause (MAP) for a period of 32 weeks (or until any ART resumption criteria were met) to assess the ability to contain viral rebound after ART interruption. Samples for the BCN02-microbiome sub-study were collected at baseline (pre-Vax), during the kick and kill intervention (after M_1_, RMD_1-2-3_ and M_2_), over MAP (from 4 to 34 weeks after ART interruption) and 24 weeks after ART resumption. Timepoints included in this sub-study are indicated by black filled triangles. **b**, 13 out of the 15 BCN02 participants were rolled over the BCN02-microbiome sub-study. * One participant (B07) not eligible for MAP due to pre-defined immune futility criteria in BCN02 trial (criteria for MAP exclusion included pVL over 2,000 copies/ml in two consecutive determinations, CD4+cell counts decrease over 50% and/or below 500 cells/mm3 and/or development of clinical symptoms suggestive of an acute retroviral syndrome) was considered a non-controller in this microbiome sub-study. Abbreviations: ART, antiretroviral therapy; MAP, monitored antiretroviral pause; pVL, plasma HIV-1 viral load; 6m, 6 months; 3yr, 3 years.

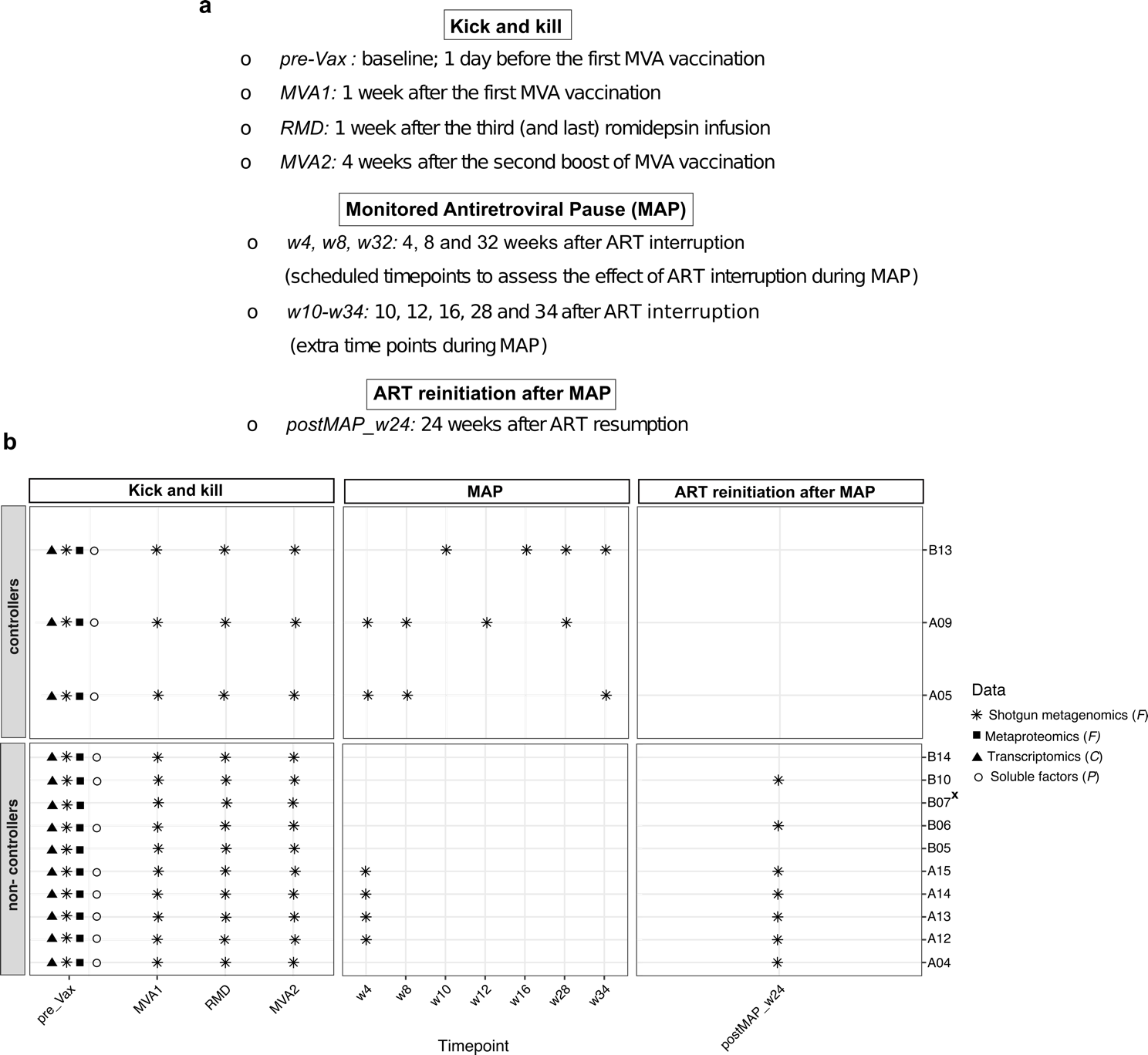

**Figure S2. Overview of sample disposition for multi-omic analysis.** **a**, Schematic representation of sampling in the BCN02-microbiome sub-study. **b**, Longitudinal sampling for multi-omics profiling including shotgun metagenomics, metaproteomics, transcriptomics and plasma proteins. In the right y-axis, participant internal identifiers are showed. The types of omics analyses performed are indicated in the legend along with the type of biological material they were performed on (C= Peripheral blood mononuclear cells (PBMCs); F=feces). (x) This participant did not enter the MAP period due to immune futility pre-defined criteria and absence beneficial HLA allele associated with natural HIV control.

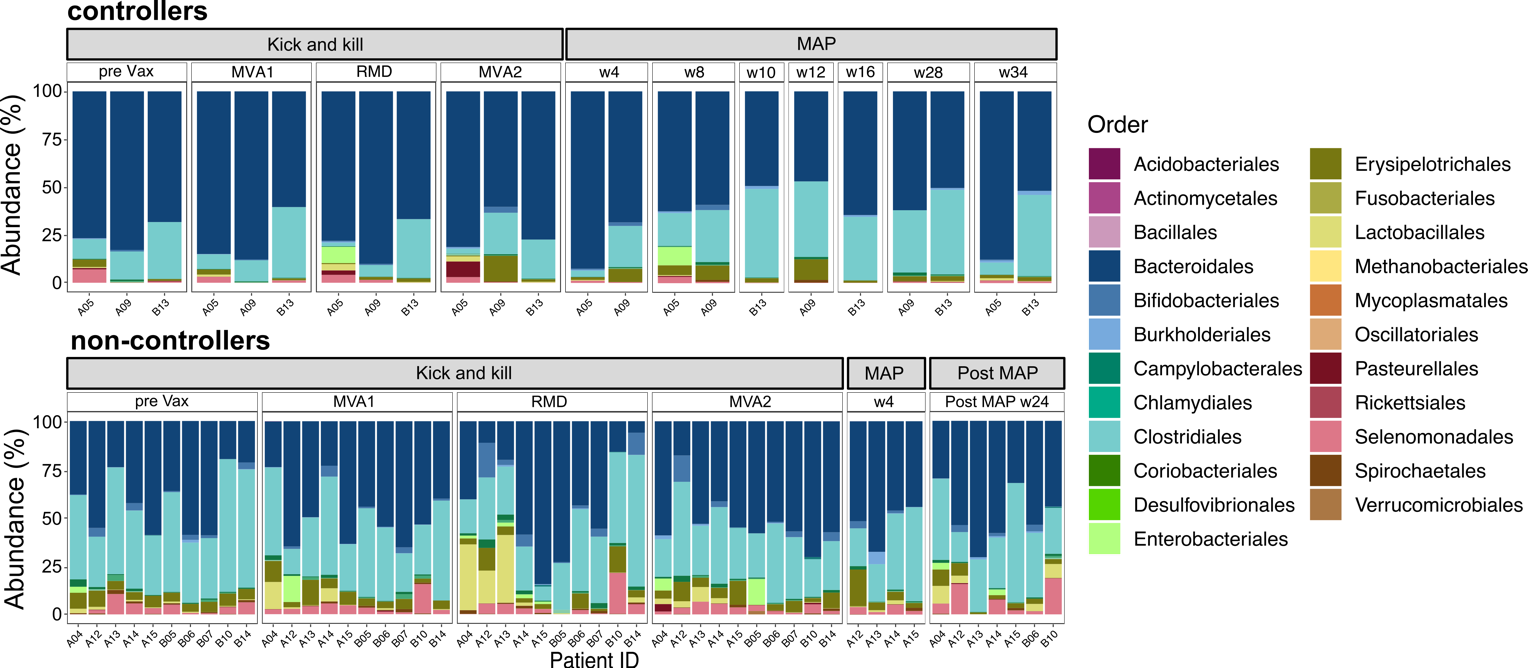

**Figure S3. Taxonomic classification of fecal samples at the order level.** Stacked bar charts showing relative abundances (in percentage) of bacterial orders at different time points for each sample within controllers and non-controllers. Each vertical bar corresponds to a study participant. Participant internal identifiers are indicated in the x-axis. Time points and phases of the trial are indicated on the top panel of each bar graph. Post MAP data are referred only to non-controllers arm. *Bacteroidales* were most represented among controllers, whereas *Clostridiales* were dominated the microbial composition in non-controllers. Abbreviations: MAP, monitored antiretroviral pause; pre_Vax, baseline (1 day before first MVA vaccination); MVA1, 1 week after first MVA vaccination; RMD, 1 week after third romidepsin infusion; MVA2, 4 weeks after second MVA vaccination.

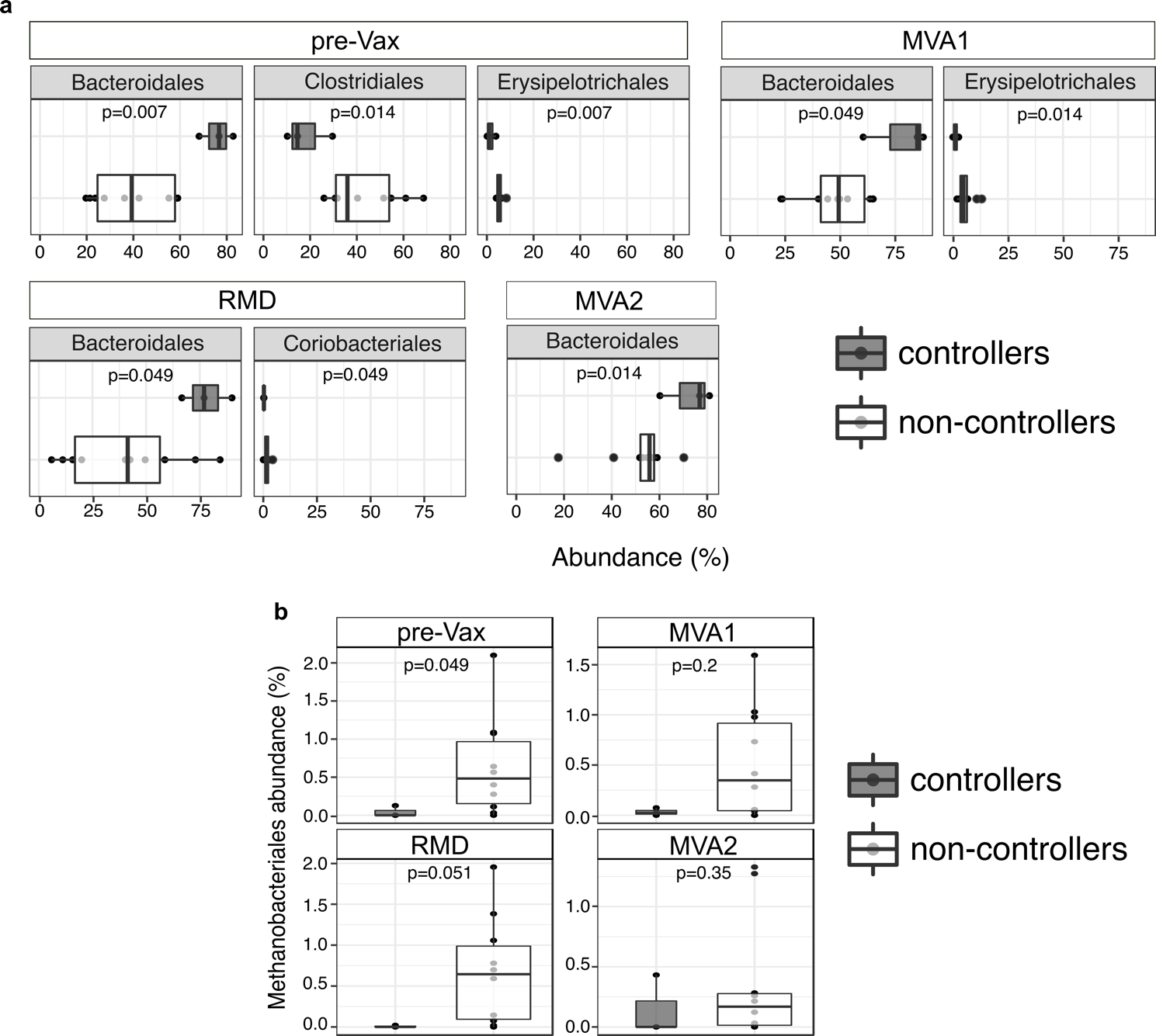

**Figure S4. Differential microbial abundance between controllers and non-controllers**. Boxplots displaying differentially abundant (**a**) bacterial and (**b**) archaeal orders at baseline (pre-Vax) and over the ‘kick and kill’ intervention (MVA1, RMD, MVA2). Relative abundances are reported as percentage on the x-axis in A. Boxes indicate the interquartile range (IQR) between the first (25^th^) and third (75^th^) quartile with the median as a vertical line inside each box. Samples and outliers are displayed with dots. Abbreviations: pre_Vax, baseline (1 day before first MVA vaccination); MVA1, 1 week after first MVA vaccination; RMD, 1 week after third romidepsin infusion; MVA2, 4 weeks after second MVA vaccination.

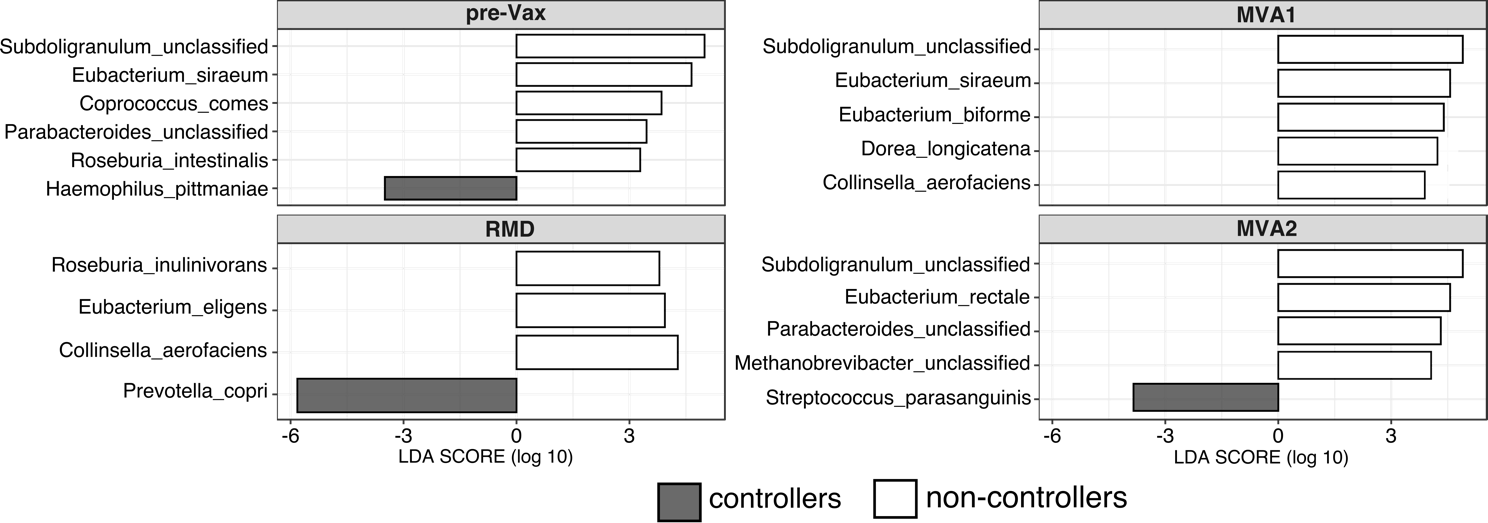

**Figure S5. Linear Discriminant Analysis (LDA) effect size (LEfSe) at the species level.** Discriminant bacterial species between controllers and non-controllers at baseline (pre-Vax) and during the ‘kick and kill’ intervention’**.** Enrichments in controllers and non-controllers are shown as grey bars with negative and white bars with positive LDA score, respectively. LDA score indicates the effect size and ranking of each discriminant feature. A p-value of < 0.05 and LDA score > 2 were considered significant in Kruskal–Wallis and pairwise Wilcoxon tests. Abbreviations: pre_Vax, baseline (1 day before first MVA vaccination); MVA1, 1 week after first MVA vaccination; RMD, 1 week after third romidepsin infusion; MVA2, 4 weeks after second MVA vaccination.

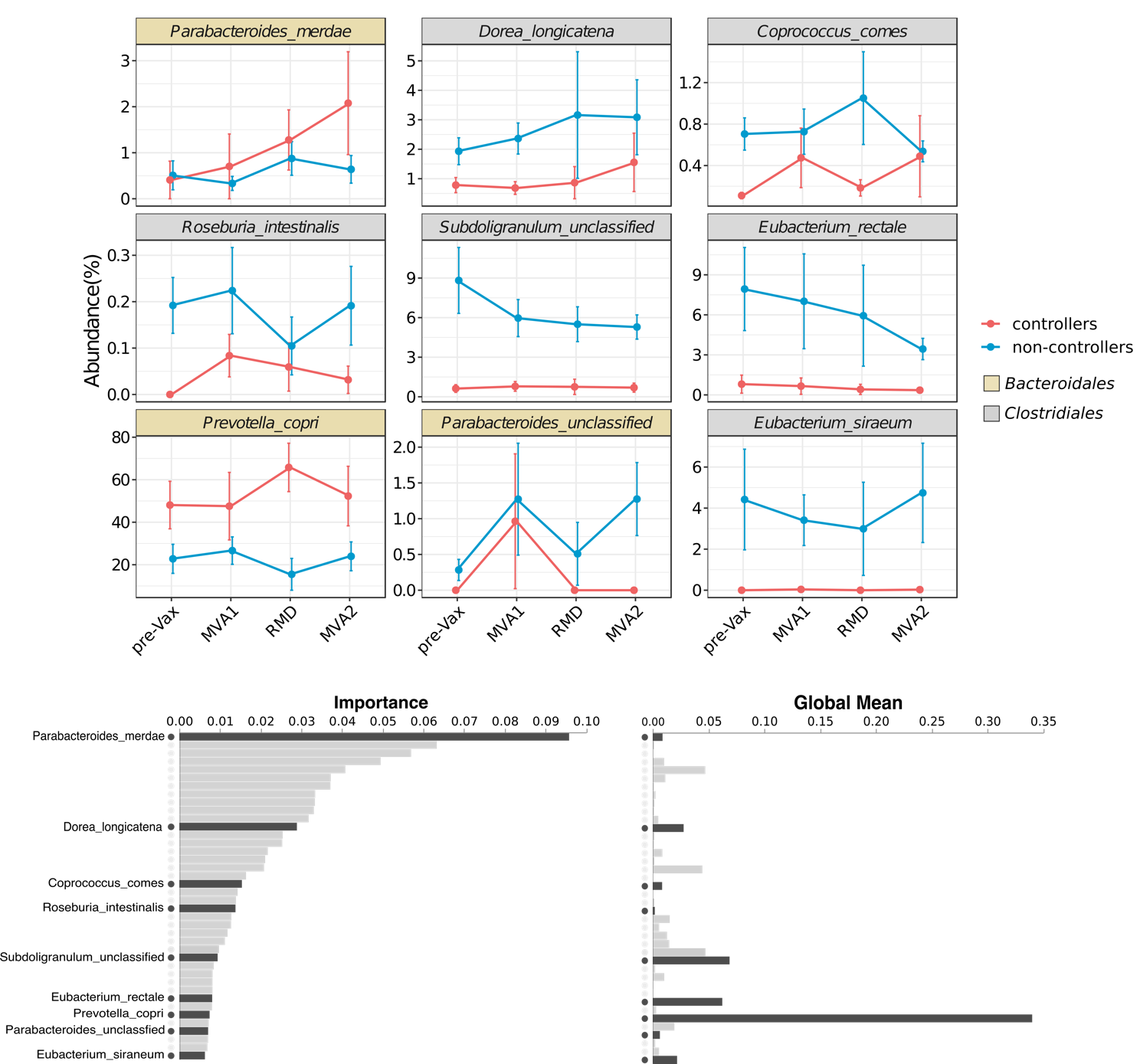

**Figure S6. Longitudinal feature-volatility analysis of bacterial species**. Feature-volatility results from q2-longitudinal analysis are plotted as volatility charts to visualize percentage relative abundance and bar charts to visualize features importance and global means. Global values from the first important feature (*Parabacteroides merdae*) and species identified by LEfSe analysis are represented for controllers (red) and non-controllers (blue) at baseline (pre-Vax) and over the ‘kick and kill’ intervention (MVA2). Solid lines represent the global group mean with standard deviations (± sd) from the mean at each time point. Dark gray horizontal bars indicate ‘Importance’ and ‘Global mean’ values for selected bacterial species. Abbreviations: pre_Vax, baseline (1 day before first MVA vaccination); MVA1, 1 week after first MVA vaccination; RMD, 1 week after third romidepsin infusion; MVA2, 4 weeks after second MVA vaccination.

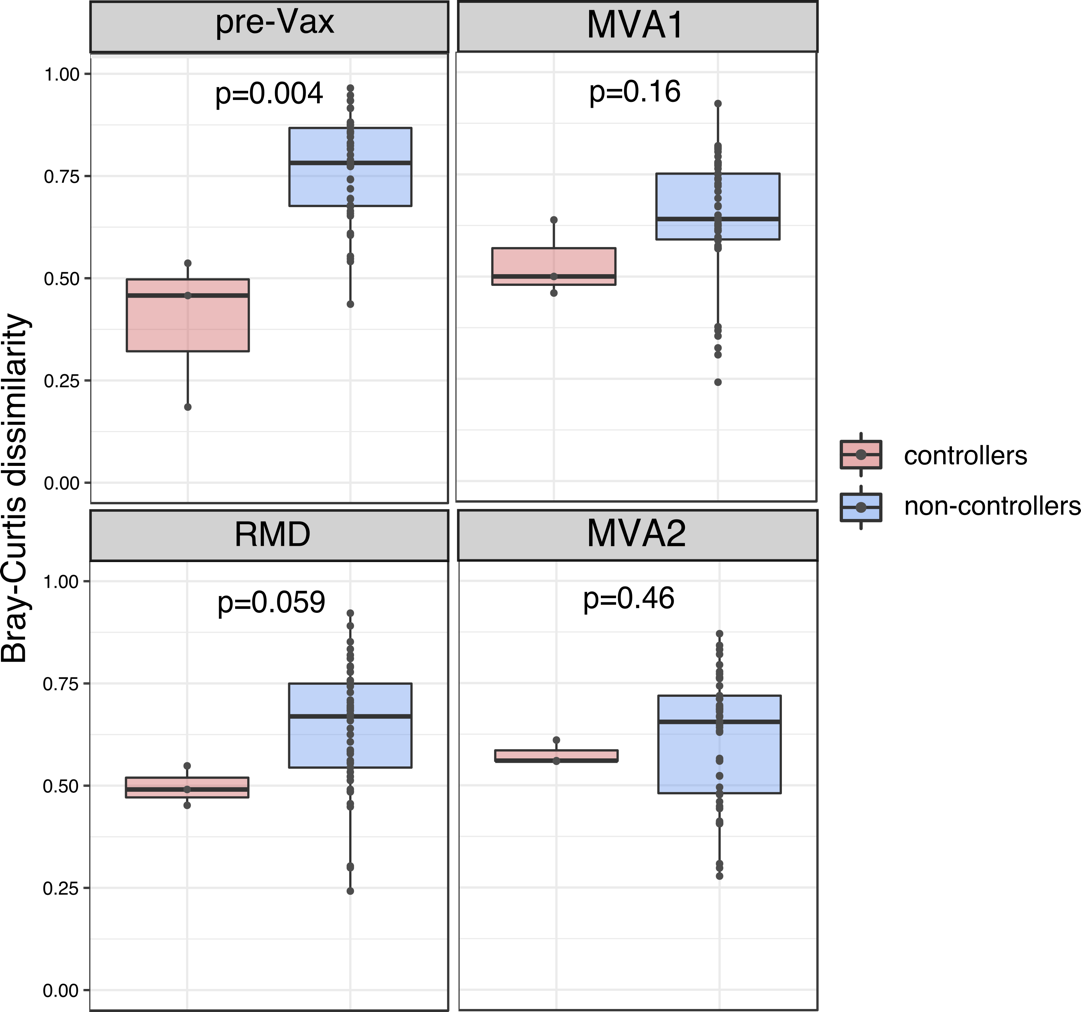

**Figure S7. Bray-Curtis dissimilarity index between controllers and non-controllers.** Comparison of Bray-Curtis dissimilarities at baseline (pre-Vax) and over the ‘kick and kill’ intervention revealed lower community dissimilarity in controllers. Boxes represent the interquartile range (IQR), the black line inside the box defines the median and whiskers represent the lowest and highest values within 1.5 IQR. Significance levels are indicated in each panel. Abbreviations: pre_Vax, baseline (1 day before first MVA vaccination); MVA1, 1 week after first MVA vaccination; RMD, 1 week after third romidepsin infusion; MVA2, 4 weeks after second MVA vaccination.

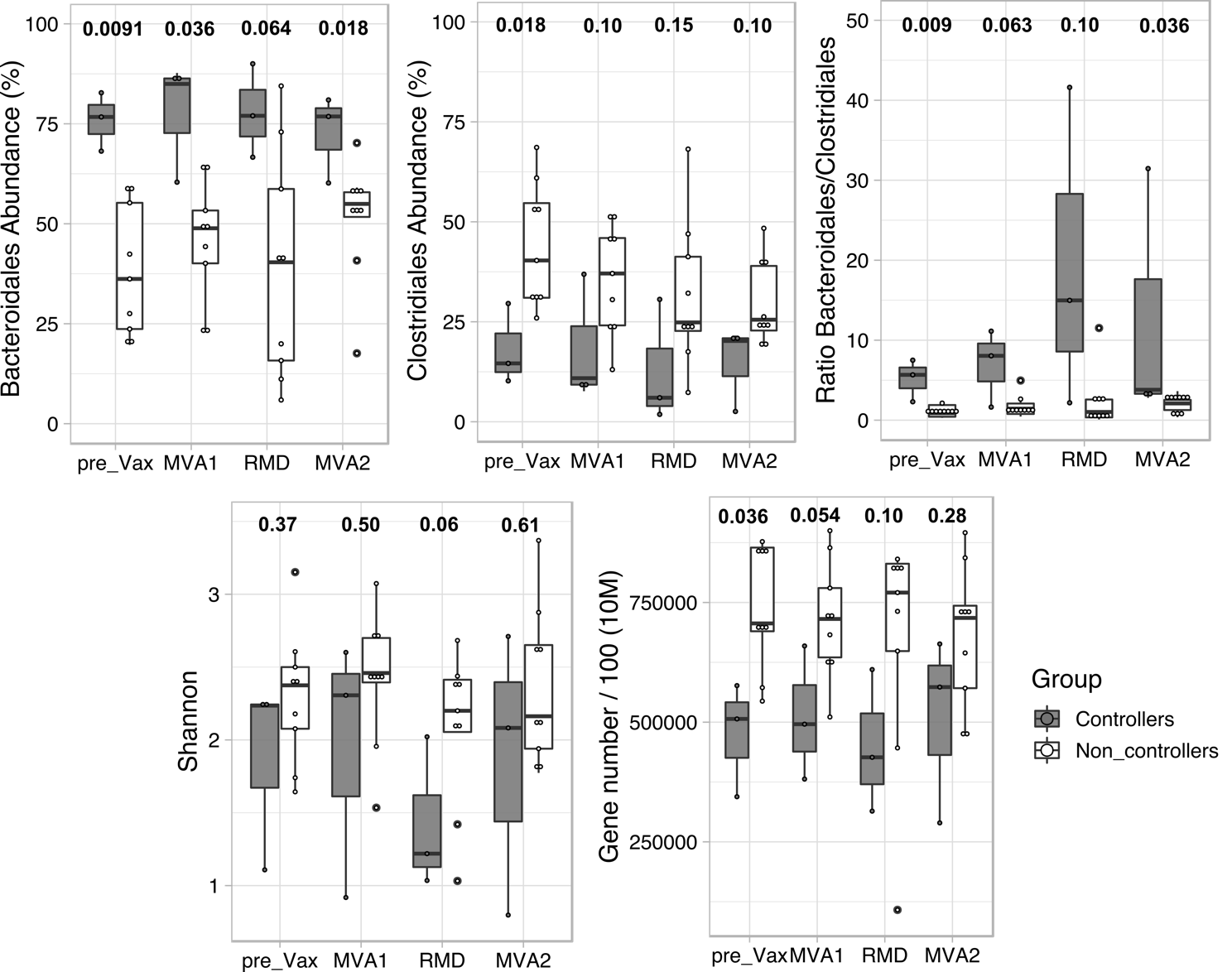

**Figure S8. Gut microbiome profiling excluding B07 participant from non-controllers arm.** Boxplots displaying longitudinal comparison (study entry and “kick and kill” intervention) of bacterial relative abundance (Bacteroidales, Clostridiales and their ratio), alpha diversity (Shannon index) and gene richness (downsampling at 10 Million reads) between controllers and non-controllers. B07 participant did not enter the MAP period due to immune futility pre-defined criteria and absence protective HLA allele associated with natural HIV control.

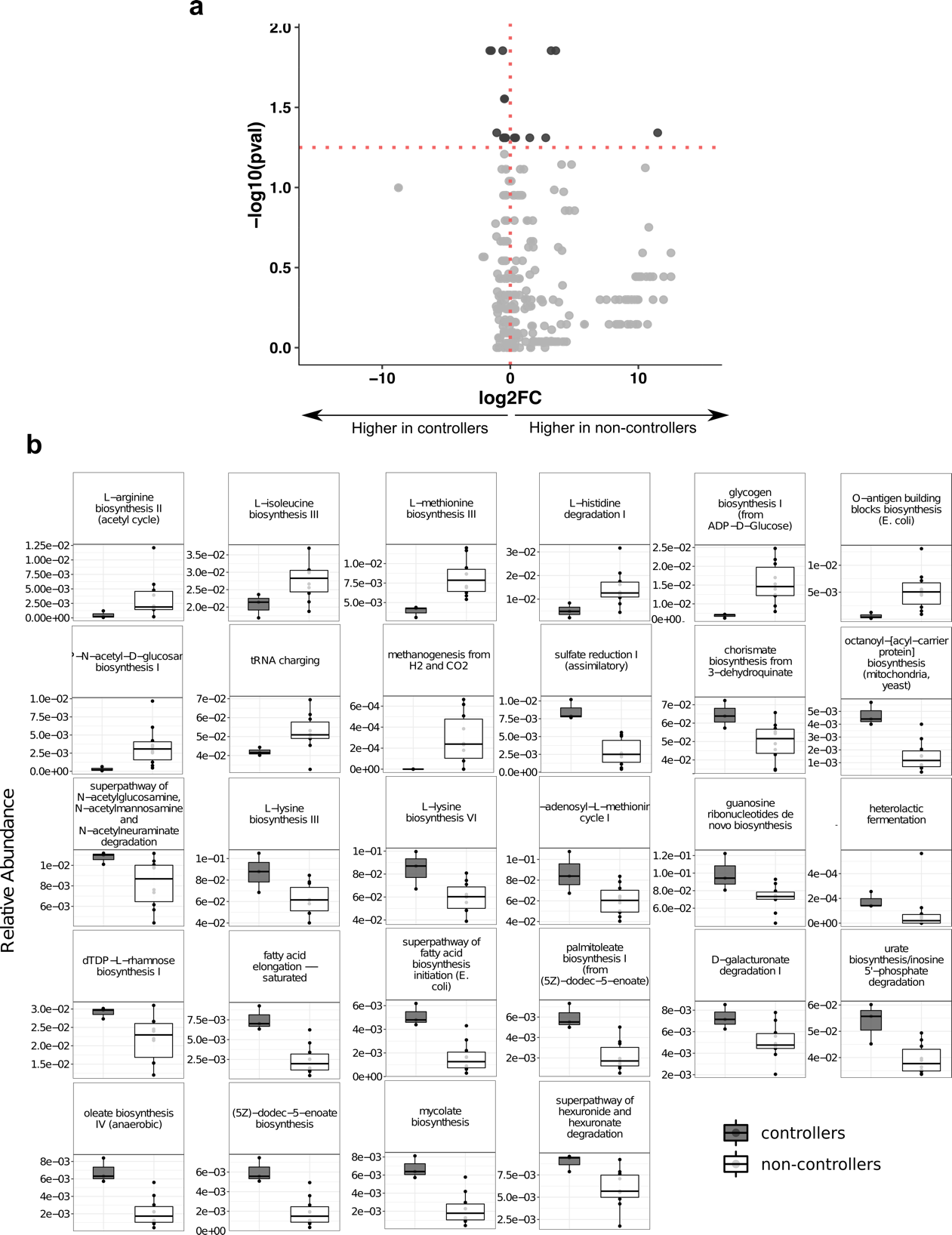

**Figure S9. Differentially abundant metabolic pathways identified at the study entry by HUMAnN2. a**, Volcano plot of microbial metabolic pathways detected in the gut microbiome of controllers and non-controllers. Log2Fold change of pathway abundance are plotted against –log10 p-value. Dotted lines in the x-axis denotes a zero-fold change, while dotted lines in the y-axis delimit a p-value value of 0.05. **b**, Boxplots showing differential metabolic pathways between controllers and non-controllers at the BCN02 study entry (*p* ≤ 0.05; Wilcoxon rank-sum test). Boxes represent the interquartile range (IQR) between the first and third quartiles (25^th^ and 75^th^ percentiles, respectively) and the black line inside each box indicate the median.

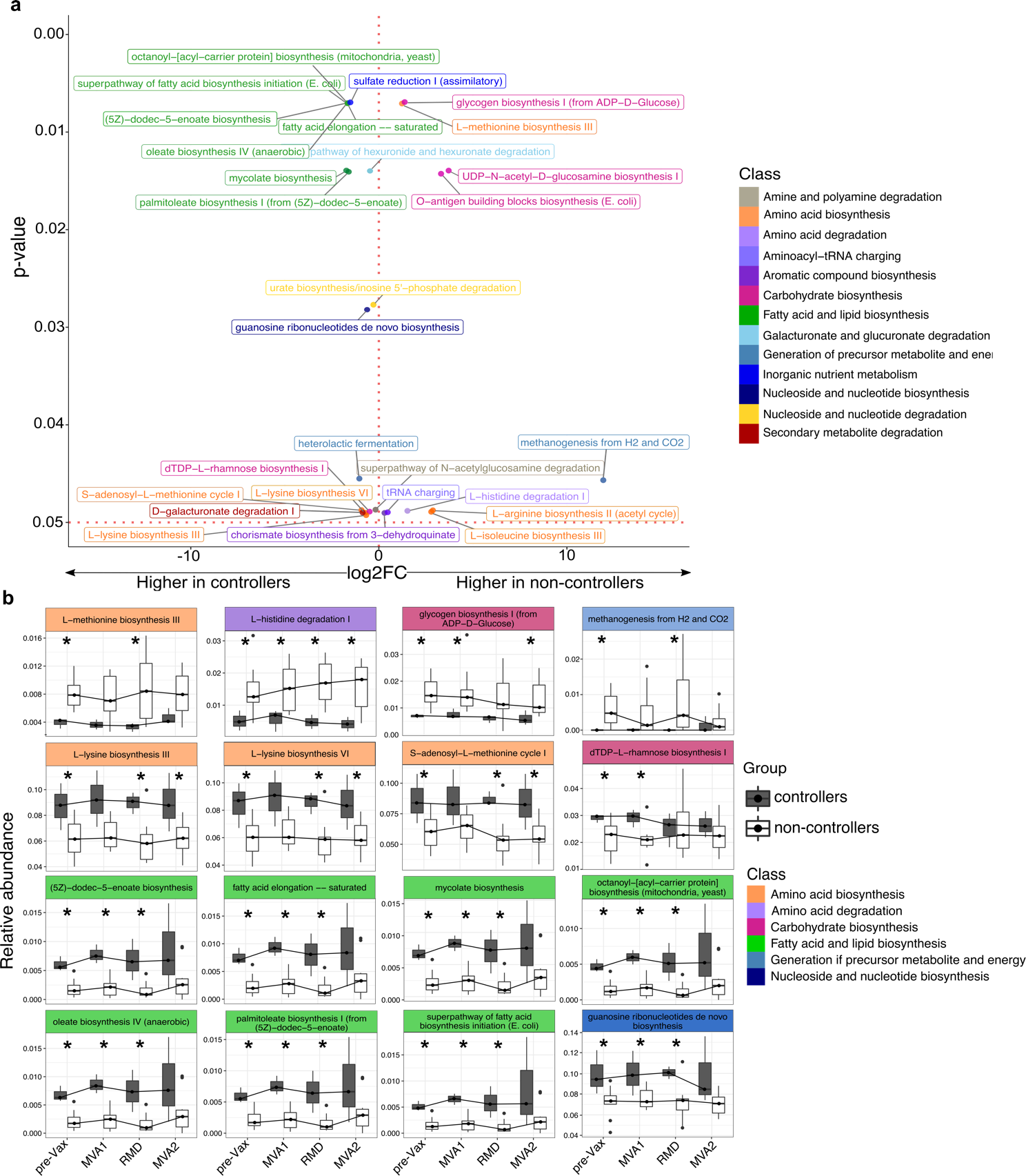

**Figure S10. Differential metabolic pathways between controllers and non-controllers. a**, Plot showing the magnitude of change of differentially abundant metabolic pathways identified at pre-Vax by HUMAnN2 and grouped by classes. Log2-transformed fold change (FC) relative abundance is plotted on the x-axis and p-values are reported on the y-axis. The red horizontal line represents the p-value cutoff at 0.05. The red vertical line delineates pathways overrepresented in non-controllers (positive FC values) and controllers (negative FC values). Functions are color-coded according to classes defined by MetaCyc. **b**, Longitudinal variation of functional pathways displaying significative differences at pre-Vax and at least one additional time point over the intervention. Black solid lines link the median value of sequential time points. Pathways are color-coded by each corresponding class. Statistical significance is highlighted by an asterisk (*P* ≤ 0.05; Wilcoxon rank-sum test, none passed multiple hypothesis testing correction). Abbreviations: pre-Vax, baseline (1 day before first MVA vaccination); MVA1, 1 week after first MVA vaccination; RMD, 1 week after third romidepsin infusion; MVA2, 4 weeks after second MVA vaccination.

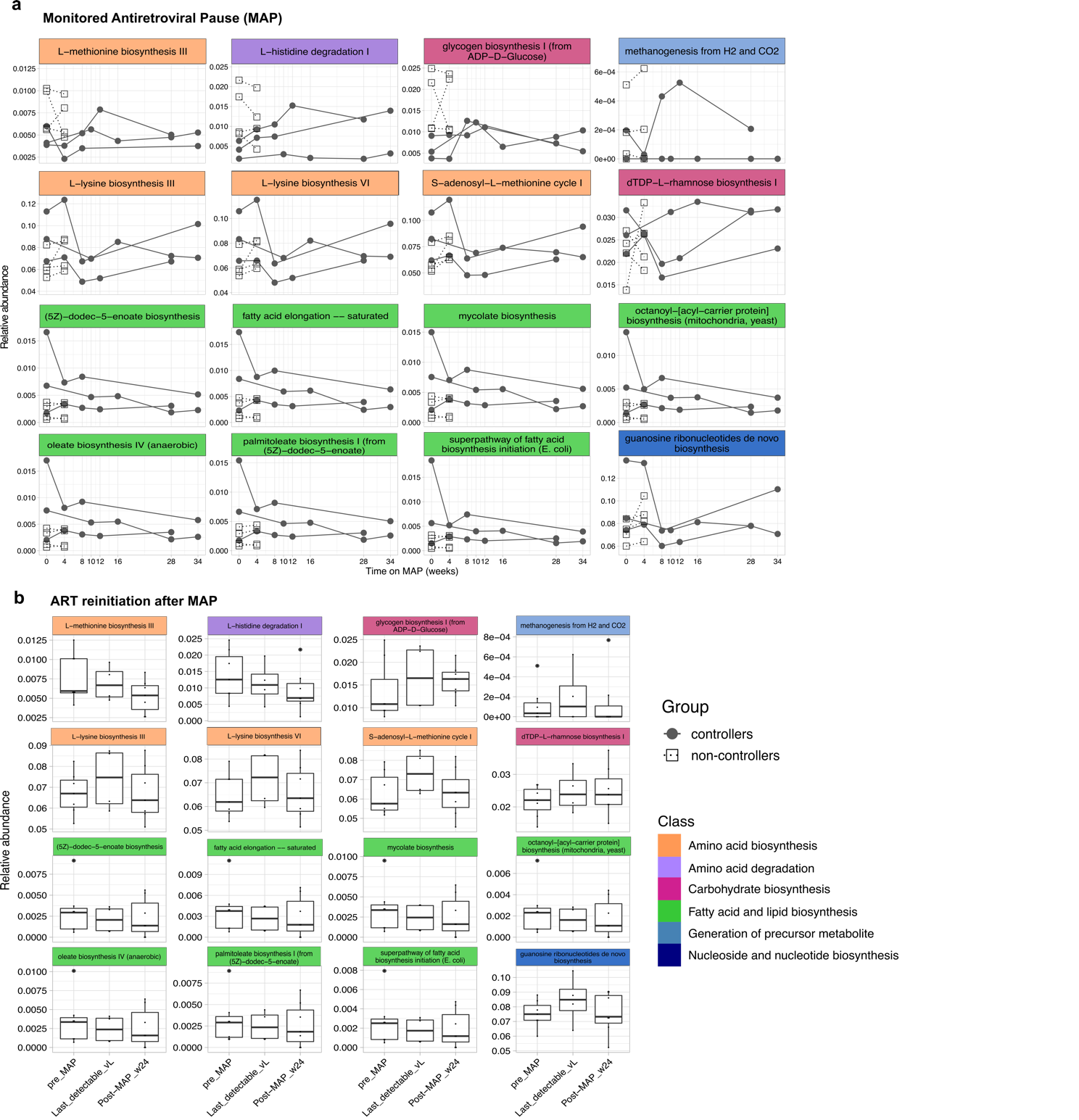

**Figure S11. Longitudinal variation of differentially abundant pathways over the trial.** Metabolic pathways showing significative differences at pre-Vax and at least one additional time point during kick and kill intervention were evaluated (**a**) during ART interruption (MAP) and (**b**) after ART reinitiation. Line plots depict pathway relative abundances for each participant belonging to controllers (grey dots) and non-controllers (white squares) arms. Boxplots illustrate pathways relative abundances in non-controllers. Each box shows the median (horizontal black line) and interquartile range between the first and third quartiles (25^th^ and 75^th^, respectively). Pathways are color-coded by each corresponding class.

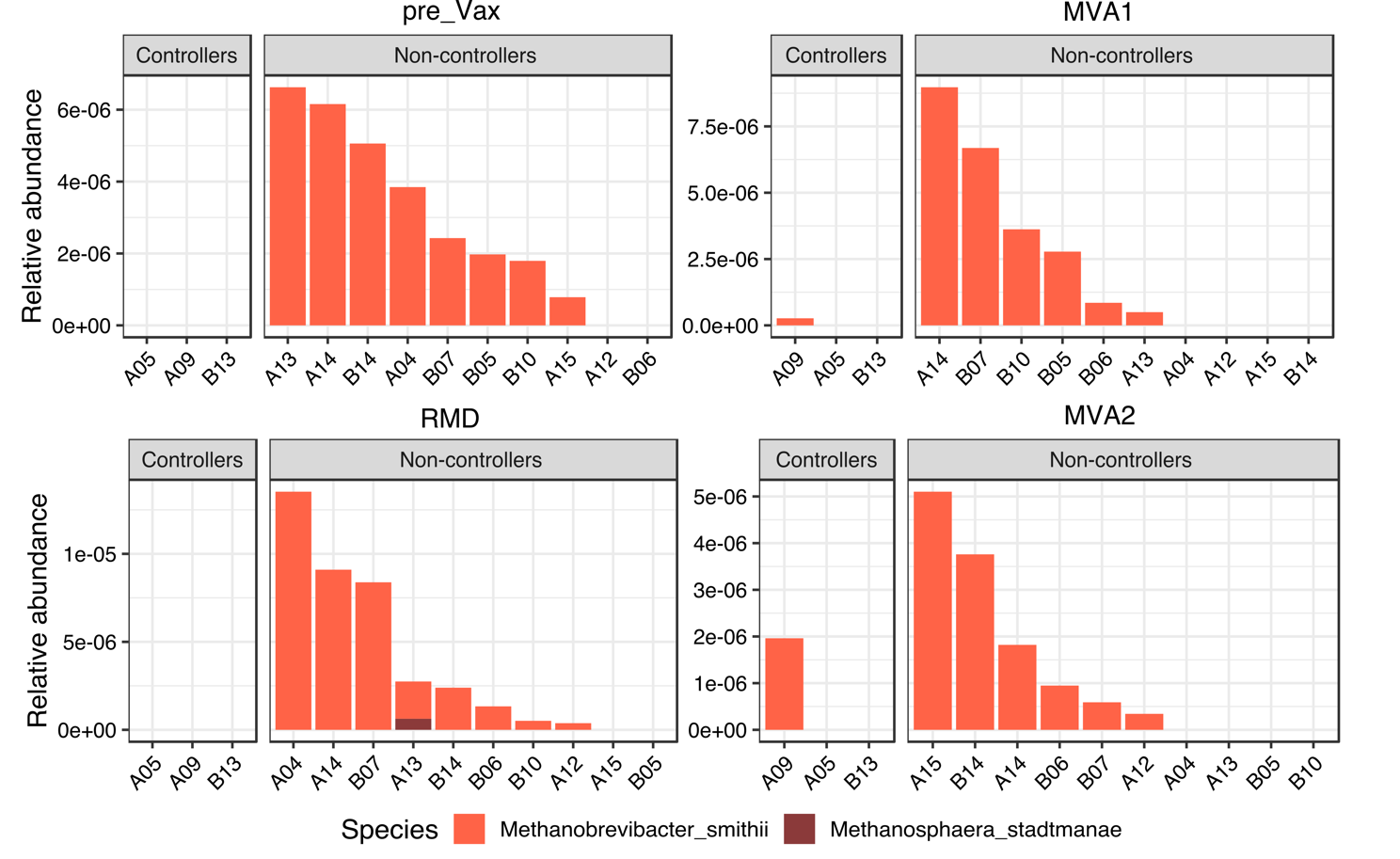

**Figure S12. Longitudinal contribution of archaeal species to the ‘methanogenesis from H_2_ and CO_2_’ pathway**. Barplots showing relative quantification of archaeal species involved in the ‘methanogenesis from H2 and CO2’ (METHANOGENESIS-PWY) pathway characterized by HUMAnN2. Abbreviations: pre_Vax, baseline (1 day before first MVA vaccination); MVA1, 1 week after first MVA vaccination; RMD, 1 week after third romidepsin infusion; MVA2, 4 weeks after second MVA vaccination.

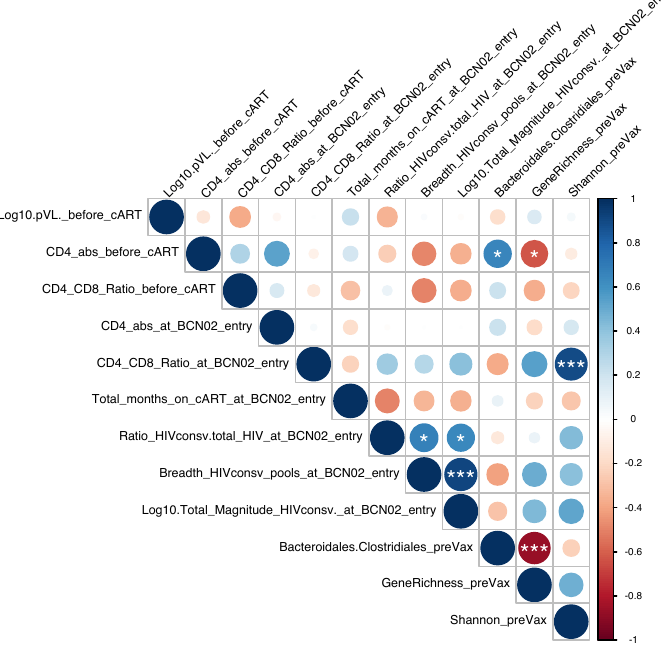

**Figure S13. Spearman’s correlation between clinical data, vaccine response and gut microbial variables.** Positive correlations are indicated in blue and negative correlations, in red. Color and size of the circles indicate the magnitude of the correlation. White asterisks indicate significant correlations (**p* < 0.05; ***p* < 0.01; ****p* < 0.001, Benjamini–Hochberg adjustment for multiple comparisons).

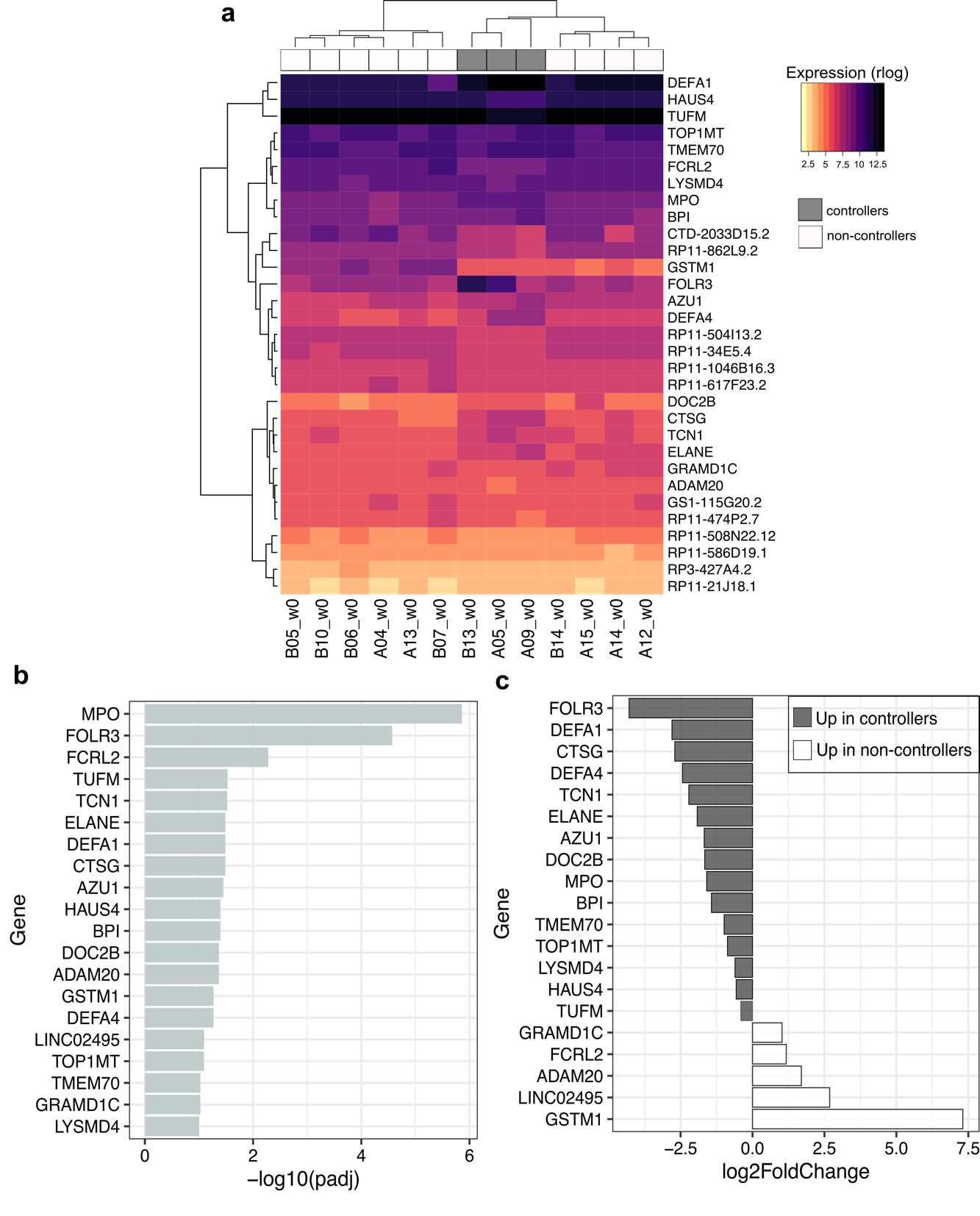

**Supplementary Figure 13. Differentially expressed PBMC host genes between controllers and non-controllers at baseline. a**, Heatmap representation of DEG (adjusted *p*-value <0.1 and log2FoldChange = 0). Gene and sample-wise hierarchical clustering was performed on individual normalized read counts (rlog). The input matrix was scaled on rows to visualize changes in expression on gene level and columns to display relatedness of samples. **c-d**,Barplots showing DEG (annotated transcripts) sorted by log10 (adjusted p-value) (**c**) and log2 (fold change) (**d**). In plot **d**, gray and white bars represent upregulated genes in controllers and non-controllers, respectively.

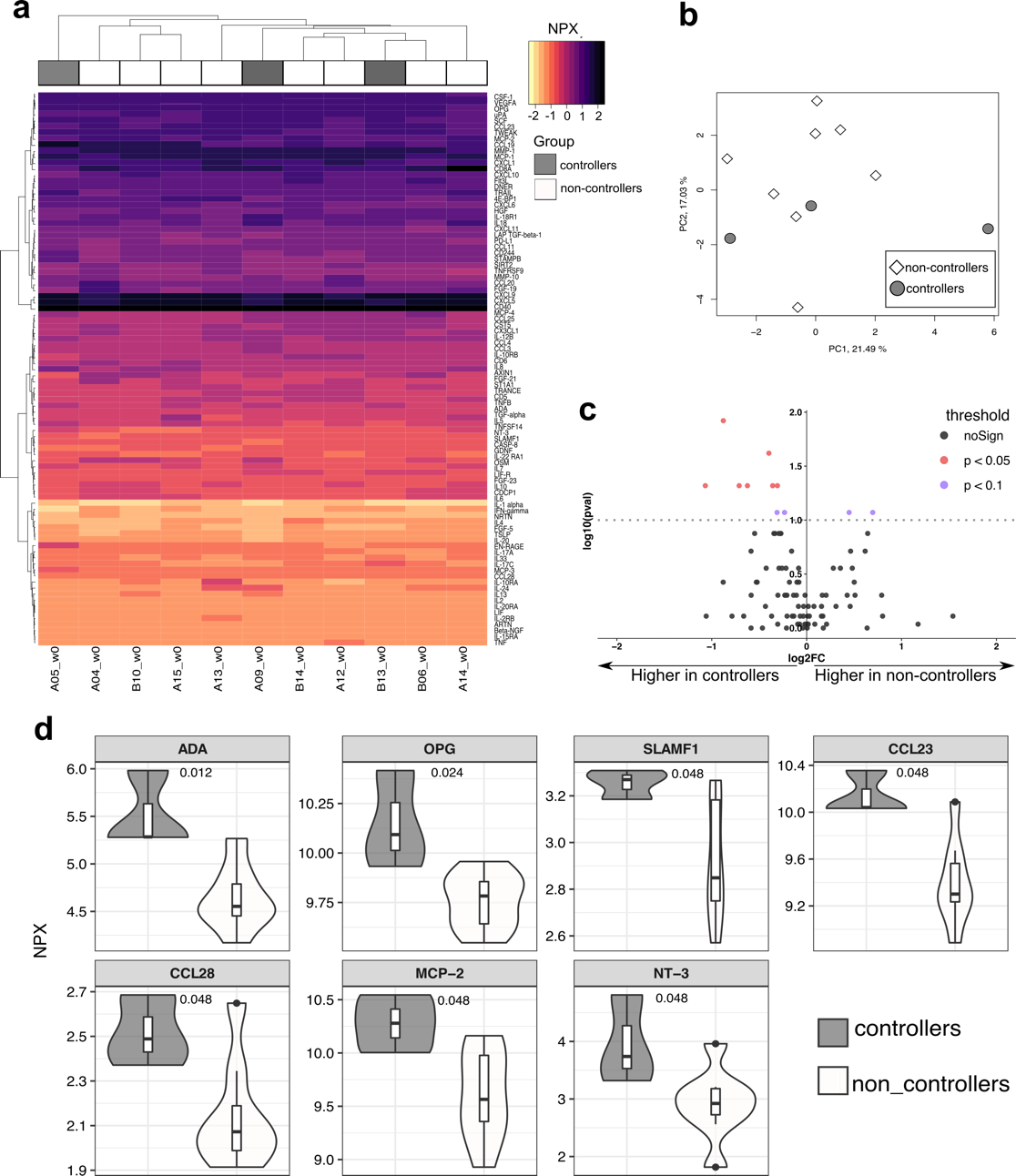

**Supplementary Figure 14. Protein inflammation markers from controllers and non-controllers at baseline.** **a**, Heatmap of plasma protein abundance based on the full Olink-inflammation panel consisting of 92 assessed proteins. Unit variance scaling was applied to NPX levels. **b**, Principal component analysis based on the full Olink-inflammation panel in controllers (gray) and non-controllers (white). The plot shows the first two principal components and their relative contribution to overall variance. **c**, Volcano plots showing differentially-expressed proteins at p-value < 0.05 (red dots), p-value < 0.1 (violet dots) and not significantly different features (black dots). Y-axis displaying the p-value and x-axis showing the fold-change in logarithmic scale. **d**, Comparative analysis of significantly different proteins (p < 0.05) between controllers and non-controllers. Y-axis indicates NPX levels for each protein.

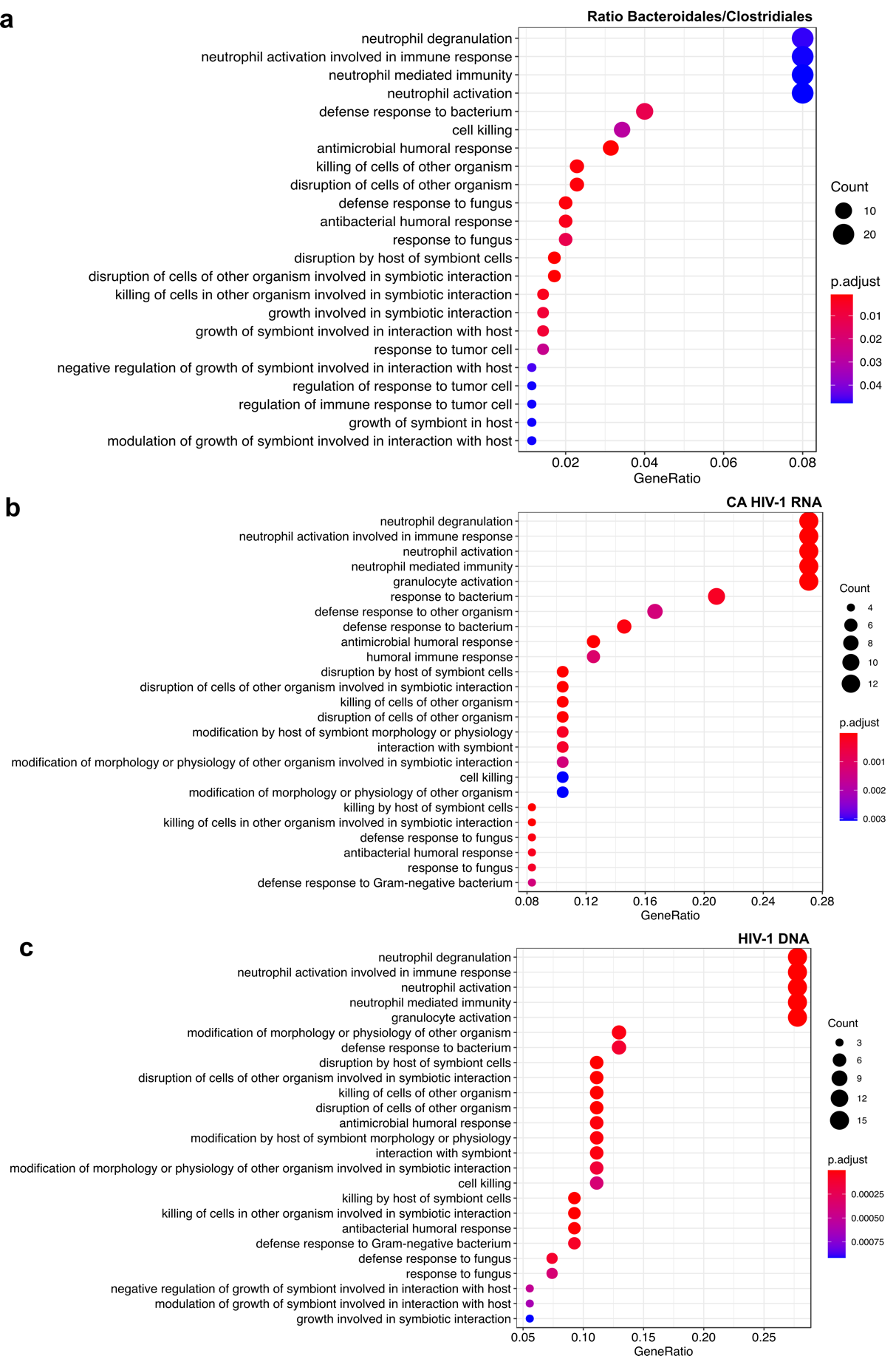

**Supplementary Figure 15. Functional enrichment of transcripts correlated with the ratio *Bacteroidales:Clostridiales* and viral reservoir.** GO enrichment of transcripts significantly correlated (q-value ≤ 0.05) with (**a**) ratio *Bacteroidales:Clostridiales*, (**b**) CA HIV-1 RNA and (**c**) HIV-1 DNA at study entry (pre-Vax) assessed by clusterProfiler. The x-axis reports the relative abundance of annotated transcripts in a given GO term, expressed as GeneRatio. Color scales indicate different thresholds of Bonferroni-adjusted p-values, and dots’ size represents the gene count of each functional GO cluster. Significantly enriched GO terms, number of genes associated to each term and adjusted *p*-values are provided in Supplementary Tables 5 and 6.

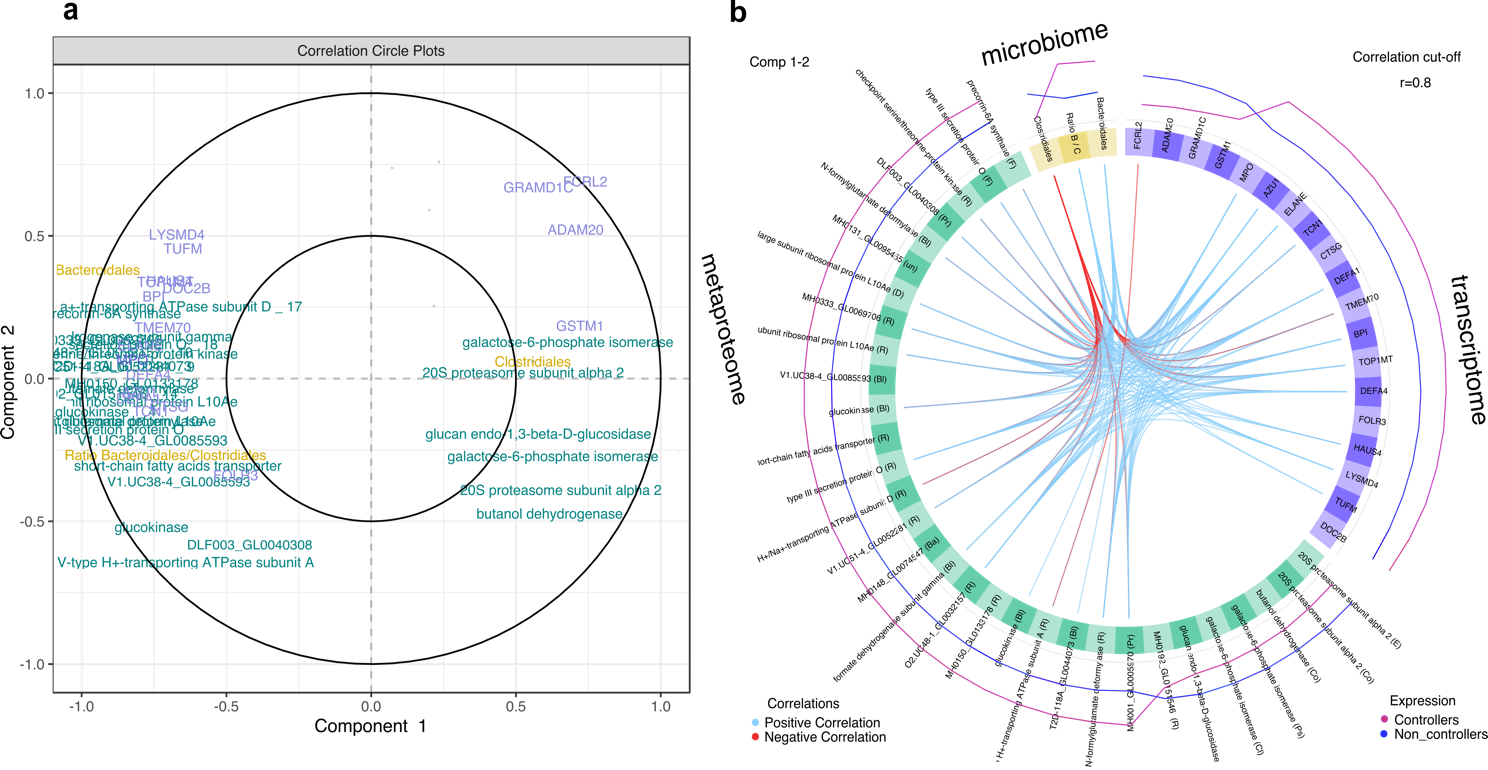

**Supplementary Figure 16. Integrated analysis of microbiome, metaproteome and transcriptome data.** Feature correlation analysis between microbial biomarkers (relative abundance), differential bacterial proteins (*p*-value ≤ 0.025) and host annotated transcripts (adjusted *p*-value ≤ 0.1 and log2 fold change = 0). (**a**) Variable plot displaying variables from each ‘-omic’ block, selected on component 1 and 2. Variable names from microbiome, metaproteome and transcriptome data are indicated in yellow, green and blue, respectively. Cluster of features indicate strong correlation between variables. (**b**) Circos plot representing correlations between variables of each ‘-omic’ block. The correlation cut-off was set at 0.8. Inner light blue and red lines correspond to positive and negative correlation between connected features, respectively. Outer purple and dark blue lines indicate the variation of each features in viremic controllers and non-controllers, respectively. Protein-associated bacterial genera are reported in parentheses. Abbreviations: Bl; *Blautia*, R; *Ruminococcus*, Pr; *Prevotella*, Ps; *Pseudoflavonifactor*, Co; *Coprococcus*, D; *Dorea*, F; *Faecalibacterium*, Ba; *Bacteroides*, E; *Eubacterium*, un; undistinguishable, Cl; *Clostridium*.
