## Additional for "Gut Microbiome Signatures Linked to HIV-1 Reservoir Size and Viremia Control": Additional Text.docx

**Additional Text for:**

**This Word file includes:**

Additional Results

Additional Methods

**Additional Results**

**Shotgun metagenomic sequencing analysis**

A median of 13 *vs* 11.4 million shotgun paired-end reads were generated for longitudinal samples belonging to controllers (n=23) and non-controllers (n=51), respectively. After trimming and filtering, 99.3% of sequencing data passed quality filters. A median of 12.9 and 11.3 non-human high-quality reads were obtained in viremic controllers and non-controllers, respectively (Supplementary Table 6). The relative abundance of microbial communities in each sample was obtained using Metaphlan2^1^. Mapped reads were mostly attributed to bacteria (99.7% controllers *vs* 97.9 % non-controllers) and smaller proportions corresponded to archaea (0.2 % controllers *vs* 0.5 % non-controllers), viruses (0.2 % controllers *vs* 1.6 % non-controllers) and eukaryotes (0.0008 % controllers *vs* 0 % non-controllers).

Consistent with previous studies of the human gut microbiota^2^, the orders *Bacteroidales* (70% in controllers and 45% in non-Controllers) and *Clostridiales* (21% in controllers and 33% in non-controllers) represented the vast majority of classified sequences in both groups, although at different proportions (Supplementary Fig. 2). *Erysipelotrichales*, *Selenomonadales*, *Lactobacillales* and *Bifidobacteriales* were detected longitudinally at lower proportions (5,3%, 4,3%, 3,2% and 2,1% in non-controllers *vs* 3.1%, 1,3%, 0,5% and 0,5% in controllers) (Supplementary Fig. 2).

**Additional Methods**

**Fecal sample collection, DNA extraction and library preparation.**

Study participants collected fecal samples at home in sterile collection tubes using standardized sampling procedures and stored immediately at approximately −20°C. Samples were then transferred to the laboratory and cryopreserved at −80°C until use. Total DNA for shotgun metagenomic sequencing was extracted and purified from each fecal sample using the PowerSoil DNA Extraction kit (MO Bio Laboratories, Carlsbad, CA), following the manufacturer’s instructions. Extracted DNA was fragmented using a Nextera-XT DNA Library Kit (Illumina, CA, USA) and one library of 300-bp average clone insert size was constructed for each sample.

**Sequencing and data quality assessment**

Metagenomic sequencing libraries were processed on an Illumina HiSeq platform (Illumina, CA, USA) by Macrogen Genomic Division. Delivered raw sequencing data were passed through the FastQC quality control software^3^. FASTQ sequence files were filtered using Trimmomatic^4^ to remove Nextera adapters and low-quality reads (minimum base quality = Q30; minimum length = 75 bp), using a sliding window set at a minimum quality of Q20 for each 30-bp-long consecutive segments. Filtered reads were then mapped against the human genome using Bowtie2^5^ to remove host DNA contamination. Reads uniquely aligned to the human reference with quality score above Q20 were discarded.

**Taxonomic classification and microbial gene richness**

Taxonomic profiles were characterized by MetaPhlAn2 software^1^ (default parameters), which used clade-specific markers to classify metagenomic reads and quantify relative abundance of taxa within each fecal sample. Microbial gene richness was assessed using the integrated gene catalog of human gut microbiome (IGC)^6^ and Bowtie2 to map filtered reads to IGC. Only filtered forward reads were used to estimate the number of unique genes and a minimum of 1 mapped read was set to consider the presence of a gene. The copy number of each gene was estimated by dividing the total reads mapping to a gene divided by the gene’s length. Gene relative abundance was measured as its copy number divided by the sum of the total gene copies in the sample, as previously reported^7^. Approximately 85% of total filtered sequences (763 835 224 reads across all samples) aligned against the IGC reference catalog (Supplementary Table 6). Based on the number of aligned sequences and rarefaction curve of new unique genes, a cut-off of 10 million mapped reads was used to compare microbial gene richness across samples.

**Microbial functional profiling**

Gene family abundance, metabolic pathway abundance and pathway coverage of each sample were determined from processed reads using HUMAnN2 (v0.11.1) with default parameters^8^. HUMAnN2 provides representative species-specific gene lists, using UniRef90^9^, MetaCyc^10^ and MinPath^11^ databases combined with MetaPhlAn2 and ChocoPhlAn pangenome databases for taxonomic identification. The resulting gene families and pathway abundance files from all samples were joined and normalized to relative abundance. The functional analysis was focused on the output of pathway abundance, which provided comprehensive quantitative insight into the functional aspects of a microbial community.

**Statistical analysis of microbiome data**

Microbial diversity and composition from shotgun metagenomic data were determined using the R (v3.5.0)^12^ packages *phyloseq*^13^, *vegan*^14^ and *ggplot2*^15^ for data visualization. Alpha diversity was estimated by the Shannon index, a measure accounting for both species abundance and evenness. Principal coordinated analysis (PCoA) based on Bray-Curtis dissimilarity was performed to display sample grouping according to their microbial composition (abundance-based). PERMANOVA (*adonis*) test based on Bray-Curtis distances was performed to estimate statistical significance of group-wise beta diversity. Normalized microbial abundances derived by MetaPhlAn2 were fed into the LEfSe algorithm^16^ to identify differentially abundant taxa between groups. A difference was considered statistically significant if LDA score >2 and *p-*value < 0.05 (Kruskal–Wallis test) after multiple test correction by FDR adjustment. To track longitudinal changes between consecutive time points, data obtained from MetaPhlAn2 were analyzed using q2-longitudinal plugin^17^. *Feature volatility* analysis was performed to assess longitudinal microbial abundance variations, comparing individual features within each group. Differences of intra-group comparison were assessed using paired Wilcoxon signed-rank test, whereas non-paired Wilcoxon signed-rank test was used for between-groups comparisons (two.sided). Unadjusted p-value ≤ 0.05 was considered statistical significance, unless otherwise specified

**Sample preparation and mass spectrometry analysis for metaproteomics**

A volume of 1.5 ml PBS was added to stool samples. Homogenization was performed by vortexing for 25 min at 4°C. A portion of the stool sample was then aliquoted and diluted further (dilution factor of 4) with PBS. Large particulates (undigested material, human cells) were pelleted out by low-speed centrifugation (300 x g for 5 min at 4°C). The supernatant was separated from the debris pellet and underwent further centrifugation (14,000 x g for 20 min at 4°C) to pellet out bacterial cells, which was then washed 3 times with PBS, vortexed, and resuspended in 250 μl of 4% SDS lysis buffer. The cells were then subjected to agitation and heat on a thermomixer (95°C for 10 min). Samples were then subjected to 3 cycles of probe sonication (1 min at 25% amplitude, 1 minute on ice), then 3 cycles of bead beating (0.1 mm silica beads). Bead beating cycles involved 60 sec of homogenization, centrifugation (14,000 x g for 5 min at RT) to pellet unlysed cells, transfer of resulting supernatant (cell lysate) into a separate tube, followed by an addition of 250 μl of lysis buffer to the unlysed cell pellet. Lysate protein content was quantified using the 2D Quant protein quantification kit (GE Healthcare Lifesciences). A trypsin digestion was carried out on 100 μg of protein via a FASP method. Urea (8 M), DTT (25 mM) and IAA (50 mM) were used for the denaturation, reduction, and alkylation of proteins, respectively. Samples were desalted using HPLC. The purified peptides were quantified using a quantitative fluorometric peptide assay (Pierce). Peptide samples were then dried down via vacuum centrifugation, and were resuspended in nano-LC buffer to a concentration of 0.25 μg/μl. An amount of 0.5 μg of peptide was injected into LC-MS for analysis. Mass spectrometry analysis of stool peptides was performed as described previously^18^, using a nano-flow Easy 1000 in line to a Orbitrap Fusion Lumos mass spectrometer (Thermo Fischer Scientific). A reference pooled stool sample was run every 10 samples to monitor MS consistency. Bacterial peptides were annotated against the human gut integrated non-redundant gene catalog (CNGdb, db.cngb.org) using the Mascot search engine (v2.4, Matrix Science), with human peptides added to limit potential homologous identifications. Search results were analyzed using Scaffold Q+ software (v4.9.0, Proteome Software). Identifications were restricted to those that passed a ≤1% FDR at the protein level, ≤0.1% FDR at the peptide level, and had ≤2 unique peptides/protein. Bacterial proteins were binned to either the order or genus level, with proteins that could not be assigned to a taxon classified as “undistinguishable”. Differences in the relative abundance of taxa were assessed using Mann-Whitney U tests, with Benjamini-Hochberg adjustment for multiple comparisons (FDR=5%). Proteins were annotated for biological functions using KEGG gene ontology.

**Isolation of PBMCs, RNA library preparation and data analysis**

The transcriptome was evaluated using RNA sequencing (RNA-seq) of peripheral blood mononuclear cells (PBMCs). Blood samples were collected prior vaccination and processed with Lymphoprep (STEMCELL technologies) We used AllPrep DNA/RNA Mini Kit kit and the Qiacube standard protocol RNAeasy Mini- Animal tissues and cells the to extract total RNA from 2M frozen PBMCs. Ribosomal RNA was removed from total RNA using RiboZero Magnetic Gold Kit and ribosomal RNA-depleted RNA from each sample was purified and fragmented. The RNASeq libraries from total RNA samples were prepared using a TruSeq™ Stranded Total RNA kit protocol (Illumina) according to manufacturer’s protocol. Each resultant library was quantified Agilent DNA 7500 Bioanalyzer assay (Agilent). The libraries were sequenced on HiSeq2000 (Illumina) in paired-end mode with a read length of 2x76bp using TruSeq SBS Kit v4 in a fraction of a sequencing v4 flow cell lane, following the manufacturer’s protocol. Image analysis, base calling and quality scoring of the run were processed using the manufacturer’s software Real Time Analysis (RTA 1.18.66.3) and followed by generation of FASTQ sequence files by CASAVA. High quality reads were then mapped to the hg19 human reference genome (GrCh38 version)^19^ using STAR v2.5.3a aligner ^20^ with ENCODE parameters. The number of reads counts aligning to each gene was estimated using RSEM v1.3.0 ^21^ and outputs from each sample were combined into a count matrix for subsequent analyses. Estimate abundances were analyzed for differentially expressed genes (DEGs) using the geometric mean method and negative binomial generalized linear models integrated in the DESeq2^22^ R package. To filter out low-expressed genes, features below a row-sum threshold of 5 were removed from the dataset. The input file contained 13 samples as columns and 58,450 genes as rows. After filtering steps, 13 samples with expression data from 27, 426 genes were obtained. Count data were transformed and normalized using DESeq2 regularized-logarithm transformation (rlog) to remove the dependence of variance on mean. A threshold of adjusted p-value < 0.1, after adjusting for multiple testing using Benjamini-Hochberg, and absolute values of log2 fold change > 0 were used to identify DEGs. Gene enrichment analysis was performed using enrichGO function from the ClusterProfiler package^23^ based on the ‘biological process’ gene ontology (GO) category. The following parameters were set as cut-off criteria: organism, *Homo sapiens*; ontology, BP; universe, filtered set of genes from the study dataset; p-value cut-off, 0.05; P-adjust method, Bonferroni; readable, T. GO enrichment results were visualized using ggplot2 and enrichGO outputs were fed into REVIGO^24^ to remove redundant GO terms and identify representative GO clusters.

**Targeted proteomic profiling of soluble factors in plasma**

Plasma samples were processed for analysis of soluble proteins. Concentrations of the protein profiles comprised in the Olink® Inflammation Panel (92 inflammation-related protein biomarkers, Olink Bioscience AB, Uppsala, Sweden)^25^ were estimated using the Proximity Extension Assay (PEA). Briefly, the multiplex immunoassay was based on protein target-specific antibodies coupled to two single-strand oligonucleotides (proximity probes) that, upon binding to their respective epitopes, generated a target sequence for a quantitative real time PCR (qRT-PCR) reaction. The Ct values from qRT-PCR were normalized by the subtraction of values for extension control, as well as an inter-plate control, and the resulting data transformed into normalized protein expression (NPX) units. A correction factor (normal background noise) was used to report arbitrary units on log2 scale, allowing relative quantification of proteins^26^. In the original dataset, data were missing for B05 and B07 participants

**‘Omic’ data correlations**

Spearman's correlation coefficients were computed for associations between gut microbial signatures, normalized individual bacterial metaproteomic, human transcriptomic data and viral reservoir size. Spearman’s rho, corresponding *p* values and p-values adjusted for multiple comparisons by the Benjamini-Hochberg method were calculated using ‘rcorr’ function within R package *hmisc*. Correlation matrices were produced using the R package *corrplot* ^27^ and adjusted p-value ≤ 0.05 was considered as a significant correlation*.* Correlation-based network analysis was conducted and visualized using *qgraph* package implemented in R^28^. Spring layout was used to classify similar terms based on the strength of their connections. Functional enrichment analysis was performed using enrichGO function within the ClusterProfiler package^23^. Correlation analysis of metagenomic, transcriptomic and metaproteomic datasets was performed using the multiblock analysis DIABLO (block.splsda function) from mixOmics R package^29^. Variable selection was based on features characterized in previous steps. A correlation threshold of 0.8 was set to determine key interactions between the selected data. Correlation networks were imported and plotted using mixOmics predefined functions. Cytoscape (v3.8.2) was used to build networks from metagenomic, transcriptomic and metaproteomic signatures^30^.

**References**

1.	Truong, D. T. *et al.* MetaPhlAn2 for enhanced metagenomic taxonomic profiling. *Nature Methods* **12,** 902–903 (2015).

2.	Arumugam, M. *et al.* Enterotypes of the human gut microbiome. *Nature* **473,** 174–180 (2011).

3.	Andrews, S., Krueger, F., Seconds-Pichon, A., Biggins, F. & Wingett, S. FastQC. A quality control tool for high throughput sequence data. Babraham Bioinformatics. *Babraham Institute* **1,** 1 (2015).

4.	Bolger, A. M., Lohse, M. & Usadel, B. Trimmomatic: a flexible trimmer for Illumina sequence data. *Bioinformatics* **30,** 2114–2120 (2014).

5.	Langmead, B. & Salzberg, S. L. Fast gapped-read alignment with Bowtie 2. *Nat Methods* **9,** 357–359 (2012).

6.	Li, J. *et al.* An integrated catalog of reference genes in the human gut microbiome. *Nat. Biotechnol.* **32,** 834–841 (2014).

7.	Le Chatelier, E. *et al.* Richness of human gut microbiome correlates with metabolic markers. *Nature* **500,** 541–546 (2013).

8.	Franzosa, E. A. *et al.* Species-level functional profiling of metagenomes and metatranscriptomes. *Nat. Methods* **15,** 962–968 (2018).

9.	Suzek, B. E., Wang, Y., Huang, H., McGarvey, P. B. & Wu, C. H. UniRef clusters: A comprehensive and scalable alternative for improving sequence similarity searches. *Bioinformatics* **31,** 926–932 (2015).

10.	Caspi, R. *et al.* The MetaCyc database of metabolic pathways and enzymes. *Nucleic Acids Res.* **46,** D633–D639 (2018).

11.	Ye, Y. & Doak, T. G. A parsimony approach to biological pathway reconstruction/inference for genomes and metagenomes. *PLoS Comput. Biol.* **5,** (2009).

12.	R Foundation for Statistical Computing. *R: a Language and Environment for Statistical Computing.* *http://www.R-project.org/* **2,** (2018).

13.	McMurdie, P. J. & Holmes, S. Phyloseq: An R Package for Reproducible Interactive Analysis and Graphics of Microbiome Census Data. *PLoS One* **8,** (2013).

14.	Oksanen, J. *et al.* vegan: Community Ecology Package. R package version 2.5-2. *Cran R* **1,** 2 (2019).

15.	Wickham, H. Package `ggplot2`: Elegant Graphics for Data Analysis. *Springer-Verlag New York* 1–222 (2016). doi:10.1093/bioinformatics/btr406

16.	Segata, N. *et al.* Metagenomic biomarker discovery and explanation. *Genome Biol.* **12,** (2011).

17.	Bokulich, N. A. *et al.* q2-longitudinal: Longitudinal and Paired-Sample Analyses of Microbiome Data. *mSystems* **3,** (2018).

18.	Klatt, N. R. *et al.* Vaginal bacteria modify HIV tenofovir microbicide efficacy in African women. *Science (80-. ).* **356,** 938–945 (2017).

19.	Pruitt, K. D., Tatusova, T. & Maglott, D. R. NCBI Reference Sequence (RefSeq): A curated non-redundant sequence database of genomes, transcripts and proteins. *Nucleic Acids Res.* **33,** (2005).

20.	Dobin, A. *et al.* STAR: ultrafast universal RNA-seq aligner. *Bioinformatics* **29,** 15–21 (2013).

21.	Li, B. & Dewey, C. N. RSEM: Accurate transcript quantification from RNA-Seq data with or without a reference genome. *BMC Bioinformatics* **12,** (2011).

22.	Love, M. I., Huber, W. & Anders, S. Moderated estimation of fold change and dispersion for RNA-seq data with DESeq2. *Genome Biol.* **15,** (2014).

23.	Yu, G., Wang, L. G., Han, Y. & He, Q. Y. ClusterProfiler: An R package for comparing biological themes among gene clusters. *Omi. A J. Integr. Biol.* **16,** 284–287 (2012).

24.	Supek, F., Bošnjak, M., Škunca, N. & Šmuc, T. Revigo summarizes and visualizes long lists of gene ontology terms. *PLoS One* **6,** (2011).

25.	Assarsson, E. *et al.* Homogenous 96-plex PEA immunoassay exhibiting high sensitivity, specificity, and excellent scalability. *PLoS One* **9,** (2014).

26.	Berggrund, M. *et al.* Protein Detection Using the Multiplexed Proximity Extension Assay (PEA) from Plasma and Vaginal Fluid Applied to the Indicating FTA Elute Micro Card^TM^. *J. Circ. Biomarkers* **5,** (2016).

27.	Wei, T. corrplot: Visualization of a correlation matrix. R package version 0.73. *URL https//github. com/taiyun/corrplot* (2013).

28.	Epskamp, S., Cramer, A. O. J., Waldorp, L. J., Schmittmann, V. D. & Borsboom, D. Qgraph: Network visualizations of relationships in psychometric data. *J. Stat. Softw.* **48,** (2012).

29.	Rohart, F., Gautier, B., Singh, A. & Lê Cao, K. A. mixOmics: An R package for ‘omics feature selection and multiple data integration. *PLoS Comput. Biol.* **13,** (2017).

30.	Shannon, P. *et al.* Cytoscape: A Software Environment for Integrated Models. *Genome Res.* **13,** 2498–2504 (2003).

**The BCN02 Study Group.** IrsiCaixa AIDS Research Institute-HIVACAT Hospital Universitari Germans Trias i Pujol, Badalona, Spain: Susana Benet, Christian Brander, Samandhy Cedeño, Bonaventura Clotet, Pep Coll, Anuska Llano, Javier Martinez-Picado, Marta Marszalek, Sara Morón-López, Beatriz Mothe, Roger Paredes, Maria C. Puertas, Miriam Rosás-Umbert, Marta Ruiz-Riol. Fundació Lluita contra la Sida, Infectious Diseases Department, Hospital Universitari Germans Trias i Pujol, Badalona, Spain: Roser Escrig, Silvia Gel, Miriam López, Cristina Miranda, José Moltó, Jose Muñoz, Nuria Perez-Alvarez, Jordi Puig, Boris Revollo, Jessica Toro. Germans Trias i Pujol Research Institute, Badalona, Spain: Ana María Barriocanal, Cristina Perez-Reche. Clinical Pharmacology Unit, Hospital Universitari Germans Trias i Pujol, Badalona, Spain: Magí Farré. Pharmacokinetic/pharmacodynamic modeling and simultation, Institut de Recerca de l’Hospital de la Santa Creu i Sant Pau-IIB Sant Pau, Barcelona, Spain: Marta Valle. Hospital Clinic- HIVACAT, IDIBAPS, University of Barcelona, Barcelona, Spain: Christian Manzardo, Juan Ambrosioni, Irene Ruiz, Cristina Rovira, Carmen Hurtado, Carmen Ligero, Emma Fernández, Sonsoles Sánchez-Palomino, and Jose M. Miró. Projecte dels NOMS-Hispanosida, BCN Checkpoint, Barcelona, Spain: Antonio Carrillo, Michael Meulbroek, Ferran Pujol and Jorge Saz. The Jenner Institute, The Nuffield Department of Medicine, University of Oxford, UK: Nicola Borthwick, Alison Crook, Edmund G. Wee and Tomáš Hanke.
